# Pharmacological activation of mitophagy prevents calcific aortic valve disease by targeting dysregulated mitochondrial-to-nuclear communication

**DOI:** 10.64898/2026.10.31.763012

**Authors:** Dao Zhou, Hanyi Dai, Junhao Qiu, Yihan Pan, Wangxing Hu, Hongyu Lu, Junhui Xue, Haochang Hu, Dilin Xu, Jingling Dai, Jin Lu, Shuangshuang Yang, Ningjing Qian, Yufei Li, Si Cheng, Ziling Mai, Jiaqi Fan, Qifeng Zhu, Yuchao Guo, Yuxin He, Jinghai Chen, Hongguang Xia, Jian’an Wang, Xianbao Liu

**Affiliations:** Department of Cardiology, The Second Affiliated Hospital, School of Medicine, Zhejiang University, Hangzhou 310009, China; State Key Laboratory of Transvascular Implantation Devices, Hangzhou 310009, China; Heart Regeneration and Repair Key Laboratory of Zhejiang province, Hangzhou 310009, China; Research Center of Clinical Pharmacy of The First Affiliated Hospital & Liangzhu Laboratory, Zhejiang University School of Medicine, Hangzhou, 311121, China; Institute of Translational Medicine, Zhejiang University School of Medicine, Hangzhou 310029, China; Department of Cardiology, Fuzhou University Affiliated Provincial Hospital; School of Medicine, Fuzhou University

## Abstract

**Background and Aims:** Calcific aortic valve disease (CAVD) lacks pharmacologic therapy. Mitophagy impairment contributes to various degenerative diseases, but whether it occurs in calcified aortic valves and during human valvular interstitial cell (hVIC) osteogenic differentiation remains unknown. We therefore investigated the role of mitophagy in CAVD and whether its pharmacological treatment prevents disease progression.

**Methods:** Single-cell RNA sequencing (scRNA-seq) was performed on human aortic valves and hVICs. TJ0113, a novel orally available mitophagy activator from our previous screening, was tested in two CAVD mouse models. Multi-omics analyses and functional rescue experiments were performed to elucidate the mechanism.

**Results:** Mitophagy was impaired in calcified human aortic valves and during hVIC osteogenic differentiation. TJ0113 activated MCL1-mediated mitophagy and attenuated disease progression in vitro and in vivo. Mechanistically, the mitophagy receptor MCL1 anchors a fraction of PRMT3 on mitochondria. During osteogenic differentiation, a dysregulated mitochondrial-to-nuclear communication is initiated by the increased release of PRMT3 into the cytosol, where it interacts with BACH1 and impairs BACH1 nuclear translocation. This leads to a widespread loss of BACH1 genomic occupancy, relieving BACH1-mediated transcriptional repression of the pro-osteogenic factor PARM1 and promoting hVIC osteogenic differentiation. Mitophagy activation by TJ0113 restored MCL1-PRMT3 binding and reversed these alterations.

**Conclusions:** Mitophagy impairment drives CAVD through dysregulated mitochondrial-to-nuclear communication mediated by the MCL1-PRMT3-BACH1-PARM1 axis. Pharmacological mitophagy activation by TJ0113 corrects this communication and attenuates disease progression, representing a promising therapeutic strategy for CAVD.

## Introduction

Calcific aortic valve disease (CAVD) is the most prevalent heart valve disease in the elderly^1, 2^. As a degenerative disease, its incidence increases rapidly with the aging of the population. According to the Global Burden of Disease study, the number of new CAVD cases worldwide increased 2.6-fold from 1990 to 2021^3^. However, no effective pharmacologic therapy is available to delay disease progression^1, 2^. Patients have no option but to wait until CAVD reaches an advanced stage and undergo aortic valve replacement. Therefore, elucidating the underlying mechanisms of CAVD and identifying potential pharmacologic therapies represent urgent and unmet clinical needs. The osteogenic differentiation of human valvular interstitial cells (hVICs), the major cellular component of the aortic valve, has been widely recognized as a critical event in CAVD progression^4, 5^. We have previously found that mitochondrial structural and functional impairments occur during hVIC osteogenic differentiation^6, 7^. This mitochondrial dysfunction, including oxidative stress imbalance, impaired biosynthesis, and dysregulated dynamics, contributes to CAVD progression^6, 7^. Indeed, emerging evidence suggests that restoring mitochondrial function represents a potential therapeutic direction^8-10^.

Mitophagy is an essential mitochondrial quality control process that selectively eliminates damaged mitochondria. Impairment of this process plays a critical role in aging and various degenerative diseases^11, 12^. In the context of CAVD, an observational study found that some mitophagy-related proteins were differentially expressed in the serum of CAVD patients, and rapamycin-induced macroautophagy was shown to reduce the calcified phenotype in diseased hVICs^13^. More recently, loss of the mitophagy receptor BNIP3 has been reported to promote hVIC osteogenic differentiation in vitro^14^. However, despite these clues, it remains unclear whether mitophagy is altered in diseased human aortic valve tissue and during hVIC osteogenic differentiation, and whether such changes play a causal role in CAVD progression.

In our single-cell RNA-sequencing (scRNA-seq) of human calcified aortic valves, we observed indications of mitophagy impairment. Of note, our group previously identified Myeloid Cell Leukemia 1 (MCL1) as a mitophagy receptor and screened a small-molecule compound, UMI-77, which selectively activates mitophagy by targeting MCL1^15, 16^. Subsequently, TJ0113, a derivative of UMI-77 with improved drug-forming properties and stronger mitophagy-inducing activity, was identified^17, 18^. Its mitophagy-activating capacity has been validated in multiple studies, and its long-term safety has been confirmed in rats and beagle dogs^19-21^. Moreover, TJ0113 reported positive top-line results from a phase II trial in early-stage Parkinson’s disease (NCT06596005) in January 2026, and advanced to a phase III trial (NCT07741045) in August 2026, highlighting its translational potential. Therefore, we conducted this study to investigate the role of mitophagy in CAVD and whether activating mitophagy could prevent disease progression.

In this study, we reveal that impaired mitophagy is a critical driver of CAVD. Mechanistically, MCL1 binds and anchors Protein Arginine Methyltransferase 3 (PRMT3) under physiological conditions. During hVIC osteogenic differentiation, MCL1-PRMT3 binding is diminished, leading to increased cytosolic PRMT3 and enhanced interaction between PRMT3 and the transcriptional repressor BTB and CNC homology 1 (BACH1), which in turn reduces BACH1 nuclear translocation and its transcriptional repression of Prostate Androgen-Regulated Mucin-like Protein 1 (*PARM1*), a pro-osteogenic factor we recently identified. This cascade constitutes a dysregulated mitochondrial-to-nuclear communication that drives hVIC osteogenic differentiation. Activating mitophagy with TJ0113 corrects this communication by restoring MCL1-PRMT3 binding, reducing PRMT3-BACH1 interaction, and facilitating BACH1 nuclear translocation and its transcriptional repression of *PARM1*. These findings demonstrate how impaired mitophagy drives CAVD and provide an effective mitophagy-activating drug with translational potential.

## Methods

Detailed methods are described in supplementary materials.

### Human samples and Ethics

Calcified aortic valves were obtained from patients who underwent surgical aortic valve replacement for severe calcific aortic valve stenosis. Patients with bicuspid aortic valve, rheumatic valvular heart disease or infective endocarditis were excluded. Control non-calcified aortic valves were obtained from patients who underwent heart transplantation, aortic dissection surgery, or surgery for aortic dilation combined with aortic valve regurgitation. All experiments involving humans were conducted in accordance with the Declaration of Helsinki and were approved by the Ethics Committee of the Second Affiliated Hospital, Zhejiang University. Written informed consent was obtained from each patient.

### Animal studies and Ethics

Specific pathogen-free (SPF) male low-density lipoprotein receptor knockout (*LDLr^−/−^*) mice on a C57BL/6J background were purchased from GemPharmatech (Nanjing, China). Mice were randomized into different groups and housed in SPF environment at a temperature of 22-24℃ with a 12-hour light/dark cycle. After acclimatization, 8-week-old mice were fed with normal chow diet (ND group), a high-fat diet (HFD group; TD-88137, SYSE Bio-tech, Changzhou, China), a high-fat diet supplemented with 0.014% TJ0113 (HFD+0.014% TJ0113 group), or a high-fat diet supplemented with 0.042% TJ0113 (HFD+0.042% TJ0113 group) for 24 weeks^21^. After 24 weeks, echocardiography was performed to evaluate aortic valve hemodynamics, and heart samples were harvested for histological evaluation^6, 7, 22^. In addition, the efficacy of TJ0113 was further assessed in another aortic valve wire injury-induced CAVD mouse model, as described in a previous study (Supplementary materials)^23^. All animal experiments were approved by the Animal Use Committee of The Second Affiliated Hospital, Zhejiang University.

### Single-cell RNA-sequencing

Human calcified and non-calcified aortic valve leaflets (one leaflet per sample, from two patients per group) were collected for scRNA-seq. Single-cell capture and barcoding were performed on the Chromium iX platform (10X Genomics). Libraries were prepared using the Chromium GEM-X Single Cell 3’ Kit (10X Genomics) and sequenced on the NovaSeq X Plus platform (PE150). Raw data were processed with Cell Ranger 8.0.1, and downstream analyses were performed using Seurat (v4.3.0.1). In addition, hVICs cultured in complete growth medium (CGM), osteogenic medium (OM), or OM supplemented with TJ0113 for four days (n=2 per group) were subjected to scRNA-seq to assess changes in cell cluster composition. Detailed methods are provided in the supplementary materials.

## Results

### Mitophagy is impaired in calcified aortic valves and during osteogenic differentiation of hVICs

Human calcified and non-calcified aortic valve leaflets were collected for droplet-based scRNA-seq using the 10X Genomics platform (Figure 1A). hVICs were found as the predominant cell population in aortic valves (Figure 1B). Kyoto Encyclopedia of Genes and Genomes (KEGG) pathway enrichment analysis of differentially expressed genes in the hVIC population between the two groups revealed significant enrichment of the mitophagy pathway (supplementary data online, Figure S1A). This enrichment was primarily driven by downregulated genes in calcified valves (Figure 1C). Further analysis revealed that multiple key genes involved in both the initiation phase (damage recognition and ubiquitin labeling) and the execution phase (autophagosomal membrane elongation and maturation) were downregulated in calcified aortic valves (Figure 1D-G, Figure S1B and S1C). Specifically, genes involved in ubiquitin labeling during ubiquitin-dependent mitophagy—including UBA52, UBC, and UBB—were significantly downregulated in calcified aortic valves, whereas BNIP3L, a key gene in the receptor-mediated mitophagy (ubiquitin-independent), was also downregulated, suggesting that recognition and tagging of damaged mitochondria were impaired in both processes (Figure 1D and 1E, Figure S1B and S1C). In addition, RABGEF1, which regulates autophagosome formation, and GABARAP, which mediates autophagosomal membrane elongation, were also downregulated, indicating that autophagosome formation for mitochondrial engulfment and clearance was compromised (Figure 1F and 1G). Collectively, these findings suggest that mitophagy is functionally impaired in hVICs from calcified human aortic valves.

**Figure 1.**
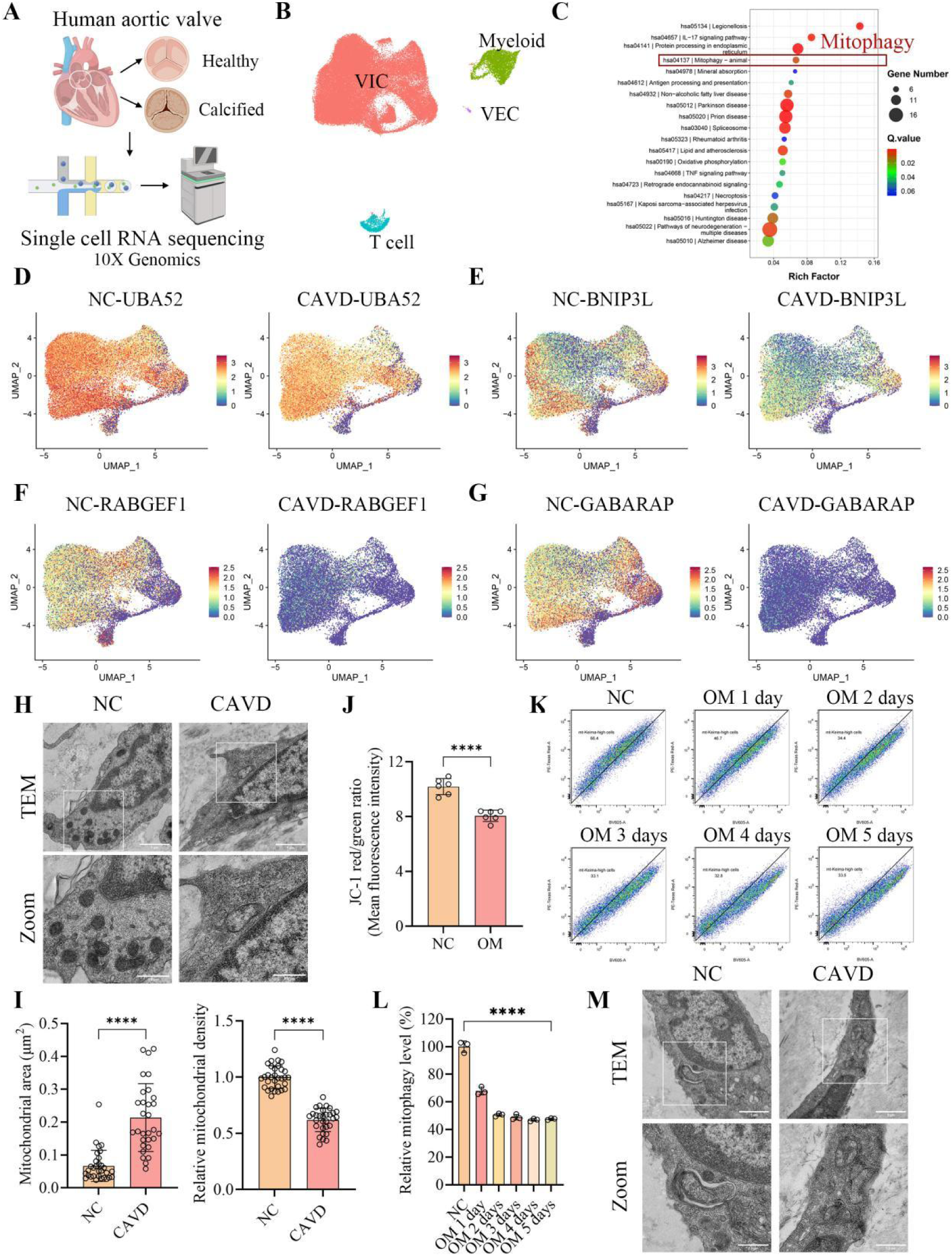
Mitophagy impairment in human calcified aortic valves and during hVIC osteogenic differentiation. (A) Schematic overview of the scRNA-seq workflow for human calcified and non-calcified aortic valve leaflets. (B) Uniform manifold approximation and projection (UMAP) of major cell types identified in human aortic valves, with VICs as the predominant population. (C) KEGG pathway enrichment analysis of downregulated genes in hVICs from calcified aortic valves, showing significant enrichment of the mitophagy pathway. (D-G) UMAP plots showing the downregulation of UBA52 (D), BNIP3L (E), RABGEF1 (F), and GABARAP (G) in hVICs from calcified aortic valves compared with non-calcified valves. (H) Representative TEM images of mitochondria in cells from calcified and non-calcified aortic valves, showing markedly swollen mitochondria with reduced electron density in calcified valves. (I) Quantification of mitochondrial area (left) and relative mitochondrial electron density (right) based on TEM images (n = 30-31 mitochondria). (J) Mitochondrial membrane potential of hVICs assessed by JC-1 staining and flow cytometry. The red/green (PE/FITC) ratio, representing the JC-1 aggregate/monomer ratio, was significantly decreased upon osteogenic differentiation (n = 6). (K) Representative flow cytometry plots of mt-Keima fluorescence in hVICs during osteogenic differentiation at different time points. (L) Quantification of relative mitophagy levels (proportion of mt-Keima-high cells relative to control) in hVICs during osteogenic differentiation at different time points (n = 3). (M) Representative TEM images showing typical mitophagic structures (autophagosomes engulfing damaged mitochondria) in non-calcified aortic valves, whereas no such structures were detected in calcified valves despite evident mitochondrial damage. For the statistical analysis, ****P < 0.0001 indicates a significant difference. Scale bars: (H, M) upper panels, 1 μm; lower panels (magnified views of boxed regions), 0.5 μm.

Given the essential role of mitophagy in mitochondrial quality control, we first examined mitochondrial ultrastructure in human aortic valve tissue. Transmission electron microscopy (TEM) revealed that mitochondria in cells from calcified aortic valves were markedly swollen and exhibited reduced electron density compared with those from non-calcified controls (Figure 1H and 1I). Consistently, mitochondrial membrane potential was significantly decreased in hVICs following osteogenic induction (Figure 1J). Logically, a decrease in mitochondrial membrane potential should trigger mitophagy. However, using mt-Keima, a well-established reporter of mitophagy^12, 24, 25^, we observed that mitophagy level was significantly reduced during osteogenic differentiation of hVICs (Figure 1K and 1L). Moreover, further TEM observation identified typical mitophagic structures (autophagosomes engulfing damaged mitochondria) in control non-calcified aortic valves (Figure 1M). In contrast, although mitochondria were clearly damaged in calcified valves, no such mitophagic structures were detected. Taken together, these findings demonstrate that mitophagy is impaired in calcified human aortic valves and during osteogenic differentiation of hVICs.

### Mitophagy activation by TJ0113 prevents calcific aortic valve disease

To investigate the functional role of mitophagy in CAVD, we first evaluated whether pharmacological activation of mitophagy could inhibit osteogenic differentiation of hVICs. Treatment with TJ0113 (5 μM, determined by concentration gradient screening; Figure S2A-C) significantly reduced osteogenic medium (OM)-induced expression of the osteogenic marker proteins ALPL and RUNX2, as well as total alkaline phosphatase activity and calcium deposition in hVICs (Figure 2A-C and Figure S2D). To further explore changes in cell cluster composition, we performed scRNA-seq on hVICs cultured under different conditions. Upon osteogenic differentiation of hVICs, cluster 4 was markedly expanded, whereas the proportion of cluster 1 was decreased (Figure 2D and 2E, Figure S2E). Notably, TJ0113 treatment reduced the proportion of cluster 4 and shifted the overall cellular landscape toward that of the control group (Figure 2E and Figure S2E).

**Figure 2.**
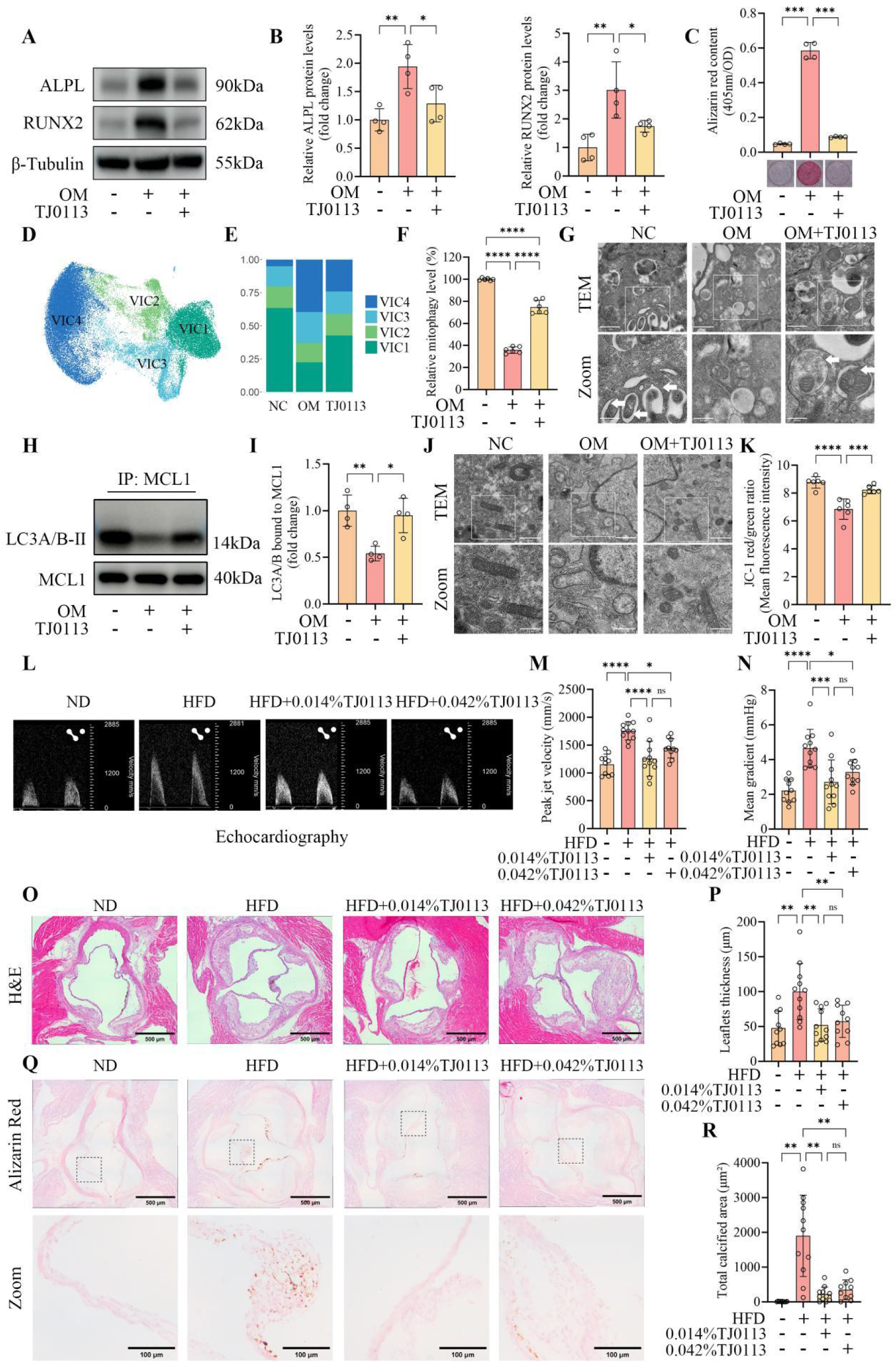
Pharmacological activation of mitophagy by TJ0113 attenuates CAVD progression in vitro and in vivo. (A) Representative Western blot images of ALPL and RUNX2 in hVICs cultured in control medium, OM, or OM supplemented with TJ0113. (B) Quantification of ALPL and RUNX2 protein levels (n = 4). (C) Alizarin Red staining and quantification of calcium deposition after 21 days of culture (n = 4). (D-E) scRNA-seq of primary hVICs cultured in control medium, OM, or OM supplemented with TJ0113. (D) UMAP plot showing four distinct subclusters of hVICs from all three groups. (E) Proportions of each hVIC subcluster across groups in one representative sample. (F) Relative mitophagy levels assessed by mt-Keima flow cytometry (n = 6). (G) Representative TEM images of hVICs cultured in control medium, OM, or OM supplemented with TJ0113, followed by bafilomycin A1 treatment for 6 hours before collection to block autophagosome-lysosome fusion. Typical mitophagic structures (white arrows) were observed in control and TJ0113-treated cells, whereas in OM-treated cells, autophagosomes were present but no mitophagic structures were detected despite evident mitochondrial damage. (H) Co-IP analysis of MCL1-LC3 interaction. hVICs cultured in control medium, OM, or OM supplemented with TJ0113 were immunoprecipitated with an anti-MCL1 antibody, and the bound LC3A/B was detected by Western blot. (I) Quantification of LC3A/B bound to MCL1, normalized to immunoprecipitated MCL1 (n = 4). (J) Representative TEM images of mitochondrial ultrastructure in hVICs. TJ0113 alleviated mitochondrial swelling, cristae loss, and vacuolization induced by osteogenic differentiation. (K) Mitochondrial membrane potential of hVICs assessed by JC-1 staining and flow cytometry (n = 6). (L) Representative pulsed-wave Doppler images of aortic valve flow velocity in *LDLr^−/−^* mice from four groups: normal chow diet (ND), high-fat diet (HFD), HFD + 0.014% TJ0113, and HFD + 0.042% TJ0113, after 24 weeks of feeding. (M) Peak jet velocity (n = 10-11). (N) Mean transvalvular pressure gradient (n = 10-11). (O) Representative H&E staining images of mouse aortic valve leaflets. (P) Quantification of aortic valve leaflet thickness (n = 10-11). (Q) Representative Alizarin Red staining images of calcific nodule deposition in mouse aortic valves. (R) Quantification of calcified area (n = 10-11). For the statistical analysis, *P < 0.05, **P < 0.01, ***P < 0.001, and ****P < 0.0001 indicate significant differences, and ns indicates no significant difference. Scale bars: (G, J) upper panels, 1 μm; lower panels (magnified views of boxed regions), 0.5 μm. (O) 500 μm. (Q) upper panels, 500 μm; lower panels (magnified views of boxed regions), 100 μm.

We next evaluated the effect of TJ0113 on mitophagy, mitochondrial structure, and function in hVICs. In hVICs expressing the mt-Keima reporter, TJ0113 treatment during osteogenic differentiation significantly increased mitophagy level (Figure 2F). To obtain direct morphological evidence, hVICs were treated with bafilomycin A1 for 6 hours to block autophagosome-lysosome fusion, followed by TEM examination^15^. Typical mitophagic structures were readily observed in control and TJ0113-treated hVICs (Figure 2G). In contrast, although autophagosomes were present in OM-treated cells, no mitophagic structures were detected despite clear mitochondrial damage (Figure 2G). Notably, while TJ0113 did not affect global macroautophagy (LC3-II expression and II/I ratio), it specifically enhanced MCL1-LC3 interaction, indicating selective activation of MCL1-mediated mitophagy (Figure 2H and 2I, Figure S3A-C).

Regarding mitochondrial structure and function, TEM showed that TJ0113 alleviated mitochondrial swelling, cristae loss, and vacuolization induced by osteogenic differentiation (Figure 2J). TJ0113 also restored the decreased mitochondrial membrane potential and reduced the elevated reactive oxygen species (ROS) levels under osteogenic conditions (Figure 2K, Figure S4A and S4B). Collectively, these results demonstrate that TJ0113 activates MCL1-mediated mitophagy and improves mitochondrial structure and function in hVICs exposed to osteogenic stimuli.

To further support the role of mitophagy in osteogenic differentiation of hVICs, we examined and knocked down PINK1, a key regulator of the ubiquitin-dependent mitophagy pathway. Its protein level was increased after osteogenic induction, consistent with the observed decrease in mitochondrial membrane potential (Figure S5A and S5B). Knockdown of PINK1 further increased ALPL expression, suggesting that inhibition of mitophagy promotes osteogenic differentiation and that the upregulation of PINK1 serves as a protective compensatory response (Figure S5A and S5C).

We next evaluated whether pharmacological activation of mitophagy prevents CAVD progression in vivo. Aortic valve stenosis was successfully established in *LDLr^−/−^* mice fed a high-fat diet for 24 weeks. Oral administration of both low-dose (0.014%) and high-dose (0.042%) TJ0113 significantly alleviated aortic valve stenosis^21^, reducing the elevated transvalvular mean gradient and peak jet velocity (Figure 2L-N). No significant differences in ejection fraction or fractional shortening were observed among the groups (Figure S6A and S6B). Histological analyses, including H&E and Alizarin red staining, revealed that TJ0113 treatment attenuated aortic valve leaflet thickening and calcific nodule deposition, respectively (Figure 2O-R). In plasma analyses, 24 weeks of TJ0113 treatment did not significantly affect metabolic parameters (including lipid profiles and blood glucose), liver function (ALT, AST), or renal function (creatinine, BUN) (Figure S7A-H). TJ0113-treated mice also had a body weight comparable to that of HFD mice (Figure S6C). In addition, the efficacy of TJ0113 was further confirmed in an aortic valve wire injury-induced CAVD mouse model (Figure S8). Similarly, both low- and high-dose TJ0113 administration alleviated aortic valve stenosis, reducing leaflet thickness and calcification (Figure S8A-C and S8F-I). Collectively, these findings demonstrate that pharmacological activation of mitophagy prevents calcific aortic valve disease both in vitro and in vivo.

### TJ0113 alleviates osteogenic differentiation of hVICs by restoring MCL1-mediated PRMT3 sequestration

Since MCL1 is a mitophagy receptor and the target of TJ0113^17, 18^, we knocked down MCL1 to verify its necessity for the function of TJ0113. MCL1 knockdown abolished the protective effect of TJ0113, leading to increased expression of ALPL and RUNX2, as well as enhanced calcium deposition in hVICs (Figure 3A-C and Figure S9A). Given that MCL1 primarily functions by binding to and sequestering other proteins^26, 27^, we performed co-immunoprecipitation (Co-IP) using an anti-MCL1 antibody followed by liquid chromatography-tandem mass spectrometry (LC-MS/MS) on hVICs from control, osteogenic differentiation, and TJ0113-treated groups. Among the candidates meeting our screening criteria, PRMT3 ranked as the top potential interacting protein, supported by reliable MS/MS spectra, and was detected in the control and TJ0113-treated groups but not in the OM group (Figure 3D and 3E). Molecular docking further predicted a high interaction probability between MCL1 and PRMT3 (docking score: -245.99, confidence score: 0.8721, Figure 3F). To validate this interaction, Co-IP experiments were performed and confirmed the endogenous binding between MCL1 and PRMT3 (Figure 3G). Importantly, this interaction was significantly weakened during hVIC osteogenic differentiation and was restored by TJ0113 treatment (Figure 3H and 3I). Notably, whole-cell lysate analysis showed that total PRMT3 protein levels remained unchanged across all conditions, suggesting that the observed changes in MCL1-PRMT3 interaction did not affect PRMT3 expression (Figure S9B-C). Given that MCL1 is a mitophagy receptor and that PRMT3 has also been reported to localize to mitochondria^28^, we investigated whether PRMT3 is anchored on mitochondria in hVICs and whether its subcellular localization changes during osteogenic differentiation. Subcellular fractionation revealed that PRMT3 was indeed present in mitochondria (Figure 3J). Moreover, consistent with the changes in MCL1-PRMT3 interaction, mitochondrial PRMT3 levels decreased while cytosolic PRMT3 levels increased during osteogenic differentiation, and these changes were reversed by TJ0113 treatment (Figure 3K and 3L). Collectively, these results indicate that MCL1 binds and anchors PRMT3, maintaining its mitochondrial localization in normal hVICs. During osteogenic differentiation, this binding is weakened, leading to the release of PRMT3 into the cytosol. Activation of mitophagy by TJ0113 reverses this process.

**Figure 3.**
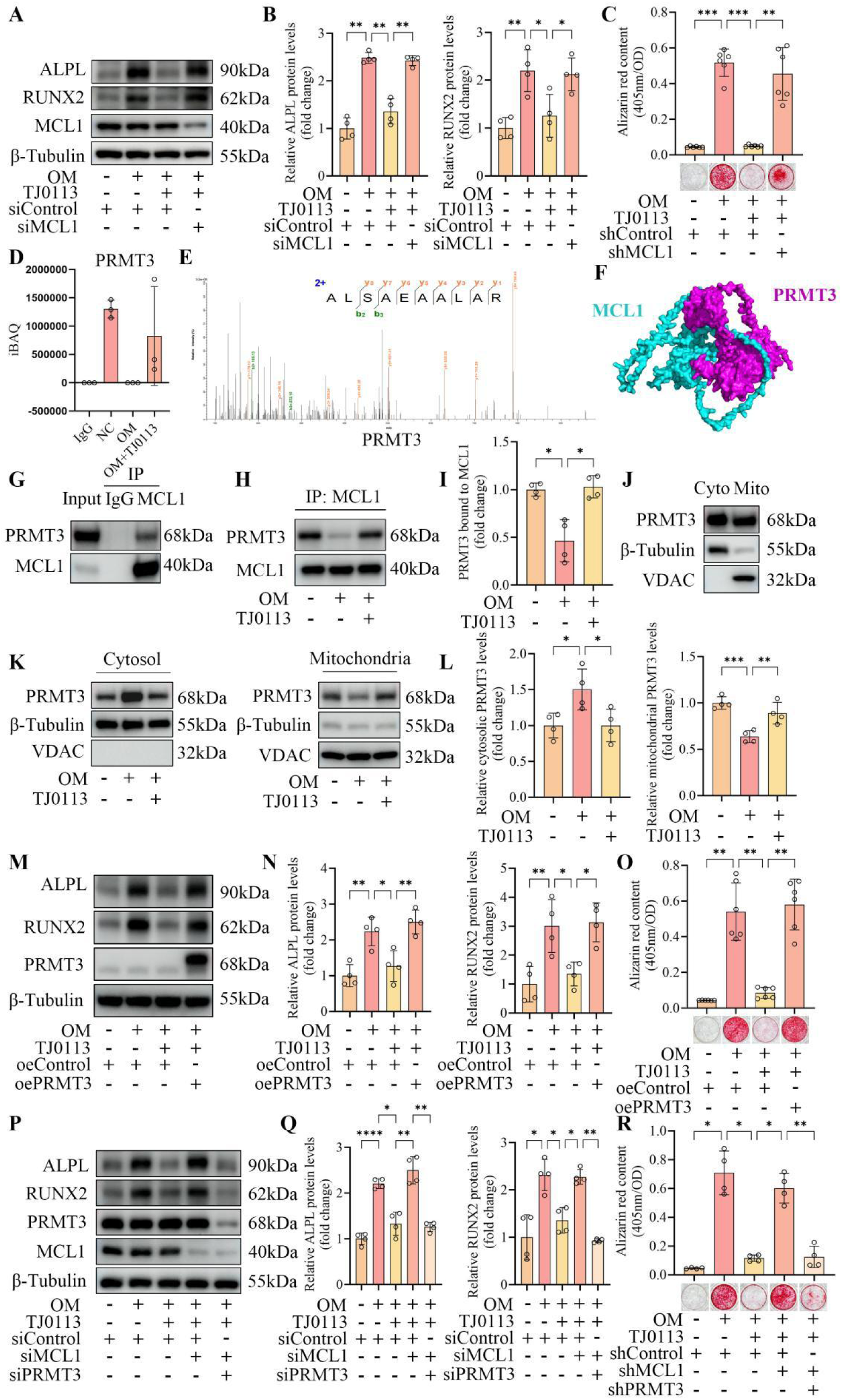
Mitophagy activation by TJ0113 restores MCL1-mediated PRMT3 sequestration on mitochondria. (A) Representative Western blot images of ALPL, RUNX2, and MCL1 in hVICs transfected with siMCL1 or siControl and cultured under different conditions. (B) Quantification of ALPL and RUNX2 protein levels (n = 4). (C) Alizarin Red staining and quantification of calcium deposition in hVICs transduced with shMCL1 or shControl lentivirus and cultured under different conditions for 21 days (n = 6). (D) iBAQ intensity of PRMT3 in hVICs from the control, OM, and OM + TJ0113 groups co-immunoprecipitated using an anti-MCL1 antibody, with an IgG control immunoprecipitated using an IgG antibody (n = 3). (E) Representative MS/MS spectrum of PRMT3 peptides identified by LC-MS/MS following Co-IP using an anti-MCL1 antibody. (F) Molecular docking model of the MCL1-PRMT3 interaction. (G) Co-IP analysis of the endogenous interaction between MCL1 and PRMT3 in hVICs. Lysates were immunoprecipitated using an anti-MCL1 antibody or an IgG control, and bound PRMT3 was detected by Western blot. (H) Representative Western blot images of PRMT3 co-immunoprecipitated using an anti-MCL1 antibody in hVICs cultured in control medium, OM, or OM supplemented with TJ0113. (I) Quantification of PRMT3 bound to MCL1, normalized to immunoprecipitated MCL1 (n = 4). (J) Western blot analysis of PRMT3 in mitochondrial and cytosolic fractions of hVICs. VDAC and β-tubulin were used as mitochondrial and cytosolic markers, respectively. (K) Representative Western blot images of PRMT3 in cytosolic (left) and mitochondrial (right) fractions of hVICs cultured under control, OM, or OM + TJ0113 conditions. (L) Quantification of cytosolic and mitochondrial PRMT3 levels, normalized to β-tubulin and VDAC, respectively (n = 4). (M) Representative Western blot images of ALPL, RUNX2, and PRMT3 in hVICs transfected with oePRMT3 or oeControl adenovirus and cultured under different conditions. (N) Quantification of ALPL and RUNX2 protein levels (n = 4). (O) Alizarin Red staining and quantification of calcium deposition in hVICs transfected with oePRMT3 or oeControl lentivirus and cultured under different conditions for 21 days (n = 6). (P) Representative Western blot images of ALPL, RUNX2, MCL1, and PRMT3 in hVICs transfected with siControl, siMCL1 + siControl (as a control for siPRMT3), or siMCL1 + siPRMT3, and cultured under different conditions. (Q) Quantification of ALPL and RUNX2 protein levels (n = 4). (R) Alizarin Red staining and quantification of calcium deposition in hVICs transfected with shControl, shMCL1 + shControl (as a control for shPRMT3), or shMCL1 + shPRMT3 lentivirus and cultured under different conditions for 21 days (n = 4). For the statistical analysis, *P < 0.05, **P < 0.01, ***P < 0.001, and ****P < 0.0001 indicate significant differences, and ns indicates no significant difference.

We next evaluated the functional role of PRMT3 and whether MCL1-mediated PRMT3 sequestration is required for the protective effect of TJ0113. PRMT3 overexpression abrogated the protective effect of TJ0113, increasing osteogenic marker expression and calcification (Figure 3M-O). Conversely, knockdown of PRMT3 significantly inhibited osteogenic marker expression and calcium nodule deposition in hVICs (Figure S10A-C). Moreover, we performed rescue experiments by knocking down PRMT3 in MCL1-knockdown hVICs. PRMT3 knockdown rescued the loss of TJ0113 efficacy caused by MCL1 knockdown, reducing ALPL and RUNX2 expression as well as calcium deposition (Figure 3P-R). Together, these results demonstrate that TJ0113 restores MCL1-PRMT3 binding, prevents PRMT3 release into the cytosol, and thereby alleviates osteogenic differentiation of hVICs.

### PRMT3 interacts with BACH1 and impairs its function, thereby promoting osteogenic differentiation

Given that PRMT3 functions via protein-protein interactions^29, 30^, we performed Co-IP with an anti-PRMT3 antibody followed by LC-MS/MS to identify its downstream effectors. BACH1 was the top candidate for PRMT3 interaction according to our screening criteria, supported by reliable MS/MS spectra (Figure 4A). Strikingly, in contrast to the MCL1-PRMT3 binding (the MS signal of PRMT3 was detected in control group and TJ0113 group but not in OM group), BACH1 was detected exclusively in the OM group (Figure 4B). Consistently, molecular docking predicted a high interaction probability between PRMT3 and BACH1 (docking score: -237.37, confidence score: 0.8516) and Co-IP experiments confirmed their endogenous interaction (Figure 4C and 4D). This interaction was markedly enhanced during hVIC osteogenic differentiation and was attenuated by TJ0113 treatment (Figure 4E and 4F). Since BACH1 protein levels remained unchanged across all conditions and previous studies have suggested that BACH1 can inhibit osteogenic differentiation of periodontal ligament cells and bone marrow mesenchymal stem cells^31, 32^, we hypothesized that enhanced PRMT3-BACH1 interaction may impair BACH1 function (Figure S11A-B). To test this, we first overexpressed BACH1 in hVICs and found that osteogenic differentiation and calcium nodule deposition were significantly inhibited (Figure 4G-I). Moreover, further BACH1 overexpression in PRMT3-overexpressing hVICs rescued the loss of TJ0113 efficacy caused by PRMT3 overexpression (Figure 4J-L). Conversely, BACH1 knockdown abolished the protective effect of either TJ0113 treatment or PRMT3 knockdown, leading to increased ALPL and RUNX2 expression in both contexts (Figure 4M-P). Together, these results indicate that during hVIC osteogenic differentiation, the released PRMT3 exhibits enhanced interaction with BACH1, thereby impairing BACH1 function. This effect promotes osteogenic differentiation and can be reversed by TJ0113 treatment.

**Figure 4.**
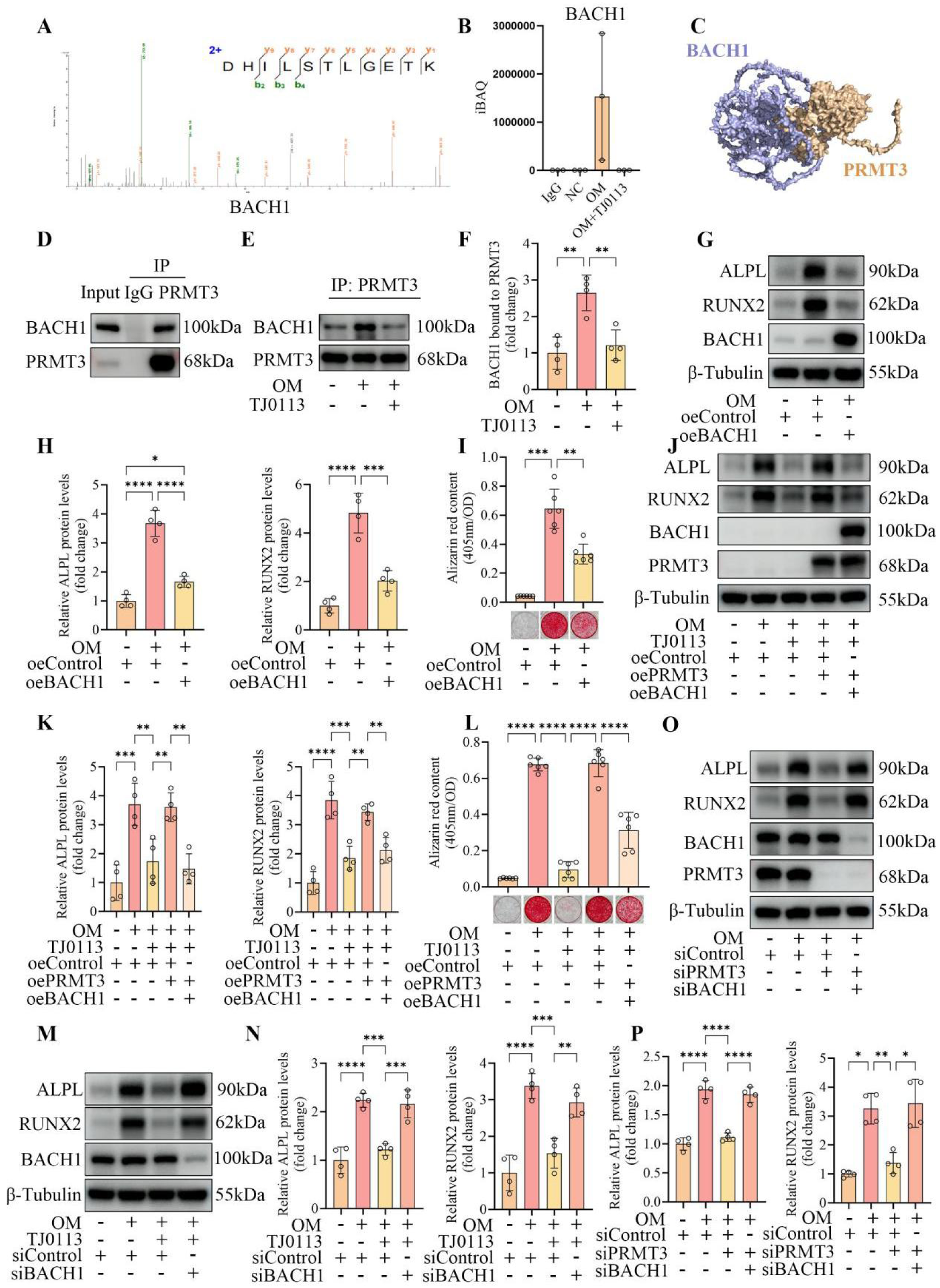
PRMT3 interacts with BACH1 and impairs its function to promote osteogenic differentiation. (A) Representative MS/MS spectrum of BACH1 peptides identified by LC-MS/MS following Co-IP using an anti-PRMT3 antibody. (B) iBAQ intensity of BACH1 in hVICs from the control, OM, and OM + TJ0113 groups co-immunoprecipitated using an anti-PRMT3 antibody, with an IgG control immunoprecipitated using an IgG antibody (n = 3). (C) Molecular docking model of the PRMT3-BACH1 interaction. (D) Co-IP analysis of the endogenous interaction between PRMT3 and BACH1 in hVICs. Lysates were immunoprecipitated using an anti-PRMT3 antibody or an IgG control, and bound BACH1 was detected by Western blot. (E) Representative Western blot images of BACH1 co-immunoprecipitated using an anti-PRMT3 antibody in hVICs cultured in control medium, OM, or OM supplemented with TJ0113. (F) Quantification of BACH1 bound to PRMT3, normalized to immunoprecipitated PRMT3 (n = 4). (G) Representative Western blot images of ALPL, RUNX2, and BACH1 in hVICs transfected with oeBACH1 or oeControl adenovirus and cultured under different conditions. (H) Quantification of ALPL and RUNX2 protein levels (n = 4). (I) Alizarin Red staining and quantification of calcium deposition in hVICs transfected with oeBACH1 or oeControl lentivirus and cultured under different conditions for 21 days (n = 6). (J) Representative Western blot images of ALPL, RUNX2, BACH1, and PRMT3 in hVICs transfected with oeControl, oePRMT3 + oeControl (as a control for oeBACH1), or oePRMT3 + oeBACH1 adenovirus, and cultured under different conditions. (K) Quantification of ALPL and RUNX2 protein levels (n = 4). (L) Alizarin Red staining and quantification of calcium deposition in hVICs transfected with oeControl, oePRMT3 + oeControl (as a control for oeBACH1), or oePRMT3 + oeBACH1 lentivirus, and cultured under different conditions for 21 days (n = 6). (M) Representative Western blot images of ALPL, RUNX2, and BACH1 in hVICs transfected with siBACH1 or siControl and cultured under different conditions. (N) Quantification of ALPL and RUNX2 protein levels (n = 4). (O) Representative Western blot images of ALPL, RUNX2, BACH1, and PRMT3 in hVICs transfected with siControl, siPRMT3 + siControl (as a control for siBACH1), or siPRMT3 + siBACH1, and cultured under different conditions. (P) Quantification of ALPL and RUNX2 protein levels (n = 4). For the statistical analysis, *P < 0.05, **P < 0.01, ***P < 0.001, and ****P < 0.0001 indicate significant differences, and ns indicates no significant difference.

### TJ0113 restores BACH1 nuclear translocation and suppresses PARM1 transcription

We next investigated how the transcriptional repressor BACH1 regulates osteogenic differentiation of hVICs. To determine whether BACH1 acts through its transcriptional regulatory function, we first performed cleavage under targets and tagmentation (CUT&Tag) sequencing on hVICs cultured under different conditions. Genome-wide binding signal analysis showed that BACH1 DNA binding signals were predominantly enriched at promoter regions, with the strongest intensity around transcription start sites (TSSs) (Figure 5A). Notably, this DNA binding was markedly reduced upon osteogenic differentiation and was restored by TJ0113 treatment (Figure 5A and 5B). Differential peak analysis identified 8,857 lost peaks and 964 gained peaks in the OM group compared with the control group. Conversely, TJ0113 treatment reversed this effect, with 7,940 gained peaks and 1,357 lost peaks compared with the OM group (Figure 5C). These results demonstrate that osteogenic differentiation induces a widespread loss of BACH1 genomic occupancy, which is effectively restored by TJ0113 treatment. Given that PRMT3 was barely detectable in the nucleus and that PRMT3-BACH1 interaction was enhanced during osteogenic differentiation, we hypothesized that this interaction may impair BACH1 nuclear translocation and subsequent DNA binding (Figure 4B-F, 5D and 5F). Nucleus and cytoplasm fractionation revealed that cytoplasmic BACH1 levels significantly increased and nuclear BACH1 levels decreased during hVIC osteogenic differentiation (Figure 5D-G). PRMT3 knockdown significantly increased BACH1 nuclear translocation (Figure 5D-G). Conversely, PRMT3 overexpression increased cytoplasmic retention of BACH1 and reduced its nuclear localization (Figure 5H-K). Moreover, TJ0113 treatment restored BACH1 nuclear translocation under osteogenic conditions (Figure 5L-O). Collectively, these results demonstrate that enhanced PRMT3-BACH1 interaction during osteogenic differentiation impairs BACH1 nuclear translocation and genomic occupancy. TJ0113 attenuates this interaction, thereby restoring BACH1 nuclear translocation and DNA binding.

**Figure 5.**
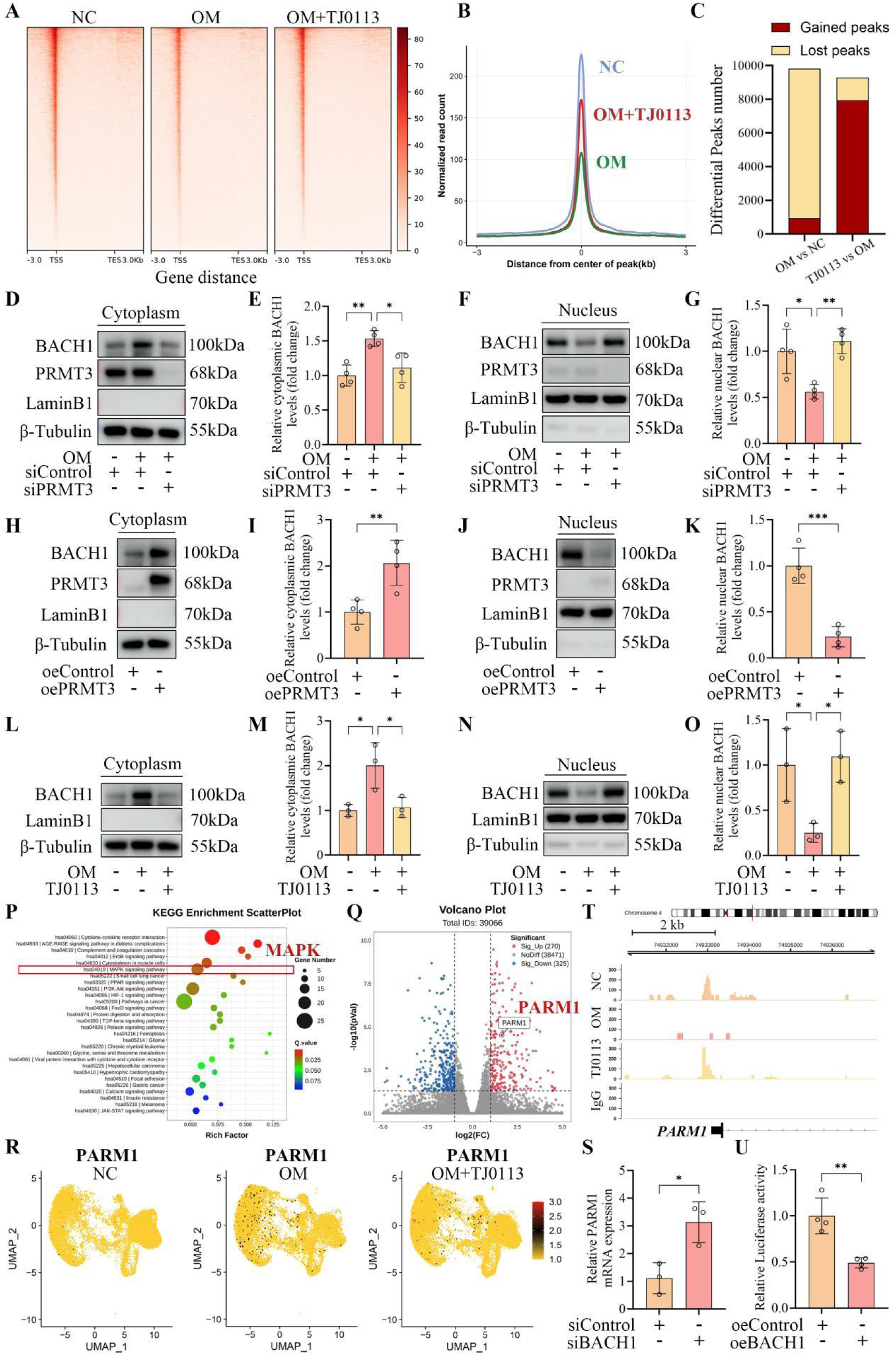
TJ0113 restores BACH1 nuclear translocation and genomic occupancy, leading to PARM1 transcriptional repression. (A) Heatmap of BACH1 binding signals across gene bodies (from 3 kb upstream of the transcription start site (TSS) to 3 kb downstream of the transcription end site (TES)) in hVICs cultured in control medium, OM, or OM supplemented with TJ0113. Genes are sorted in descending order by signal intensity. (B) Average BACH1 binding signal profiles across all peaks in the three groups. (C) Numbers of lost and gained BACH1 binding peaks in the OM group versus the control group and in the OM + TJ0113 group versus the OM group. (D) Representative Western blot images of BACH1 and PRMT3 in the cytoplasmic fraction of hVICs with or without PRMT3 knockdown under different culture conditions. (E) Quantification of cytoplasmic BACH1 levels, normalized to β-tubulin (n = 4). (F) Representative Western blot images of BACH1 and PRMT3 in the corresponding nuclear fraction. (G) Quantification of nuclear BACH1 levels, normalized to Lamin B1 (n = 4). (H) Representative Western blot images of BACH1 and PRMT3 in the cytoplasmic fraction of hVICs transfected with oePRMT3 or oeControl. (I) Quantification of cytoplasmic BACH1 levels, normalized to β-tubulin (n = 4). (J) Representative Western blot images of BACH1 and PRMT3 in the corresponding nuclear fraction. (K) Quantification of nuclear BACH1 levels, normalized to Lamin B1 (n = 4). (L) Representative Western blot images of BACH1 in the cytoplasmic fraction of hVICs cultured in control medium, OM, or OM supplemented with TJ0113. (M) Quantification of cytoplasmic BACH1 levels, normalized to β-tubulin (n = 3). (N) Representative Western blot images of BACH1 in the corresponding nuclear fraction. (O) Quantification of nuclear BACH1 levels, normalized to Lamin B1 (n = 3). (P) KEGG pathway enrichment analysis of differentially expressed genes in BACH1-knockdown hVICs compared with control hVICs, showing significant enrichment of MAPK signaling. (Q) Volcano plot of differentially expressed genes in BACH1-knockdown hVICs compared with control hVICs. PARM1 was significantly upregulated upon BACH1 knockdown. (R) UMAP plots showing PARM1 expression in hVICs cultured in control medium, OM, or OM supplemented with TJ0113. PARM1 expression was increased in the OM group and reduced in the OM + TJ0113 group. (S) RT-qPCR analysis of PARM1 mRNA levels in BACH1-knockdown and control hVICs (n = 3). (T) Genome browser tracks showing BACH1 binding signals at the PARM1 promoter in hVICs cultured in control medium, OM, or OM supplemented with TJ0113, with an IgG control. BACH1 binding peaks at the PARM1 promoter were detected in the control and OM + TJ0113 groups but were absent in the OM group. (U) Dual-luciferase reporter assay of PARM1 promoter activity in HEK293T transfected with BACH1 overexpression or control plasmids (n = 4). Luciferase activity was normalized to Renilla luciferase. For the statistical analysis, *P < 0.05, **P < 0.01, and ***P < 0.001 indicate significant differences.

To identify downstream targets of BACH1, we performed bulk RNA-seq on BACH1-knockdown and control hVICs. Among KEGG enriched pathways and the differentially expressed genes, we noticed that MAPK signaling was significantly enriched and that PARM1, a pro-osteogenic factor previously identified by our group as a CAVD driver through MAPK signaling^22^, was significantly upregulated upon BACH1 knockdown (Figure 5P and 5Q). Further analysis of our scRNA-seq data showed that TJ0113 treatment reduced PARM1 mRNA expression in hVICs (Figure 5R). Real-time quantitative polymerase chain reaction (RT-qPCR) confirmed that BACH1 knockdown significantly increased PARM1 mRNA levels (Figure 5S). To determine how BACH1 regulates PARM1 transcription, we analyzed our CUT&Tag sequencing data and detected a BACH1 binding peak at the PARM1 promoter (Figure 5T). This peak was lost upon hVIC osteogenic differentiation and was restored by TJ0113 treatment (Figure 5T). Dual-luciferase reporter assays further confirmed that BACH1 overexpression suppressed PARM1 promoter activity (Figure 5U). At the protein level, PARM1 expression was reduced by either TJ0113 treatment or PRMT3 knockdown under osteogenic conditions (Figure S12A-D). Importantly, additional BACH1 knockdown reversed this suppression in both settings (Figure 6A-D). In overexpression experiments, BACH1 overexpression reduced PARM1 levels (Figure 6E and 6F). Moreover, PRMT3 overexpression increased PARM1 expression in TJ0113-treated hVICs, and additional BACH1 overexpression again reduced PARM1 expression (Figure 6G and 6H). Consistently, in CAVD mouse models, TJ0113 administration significantly reduced PARM1 expression in aortic valves (Figure 6I and 6J, Figure S13A and S13B). Finally, we evaluated whether the protective effects of TJ0113 and BACH1 are mediated through PARM1 suppression. PARM1 overexpression abrogated these protective effects, leading to increased osteogenic marker expression and calcium deposition (Figure 6K-P). Collectively, these results demonstrate that TJ0113 restores BACH1 nuclear translocation and binding to the PARM1 promoter by attenuating PRMT3-BACH1 interaction, thereby suppressing PARM1 transcription and ultimately alleviating hVIC osteogenic differentiation.

**Figure 6.**
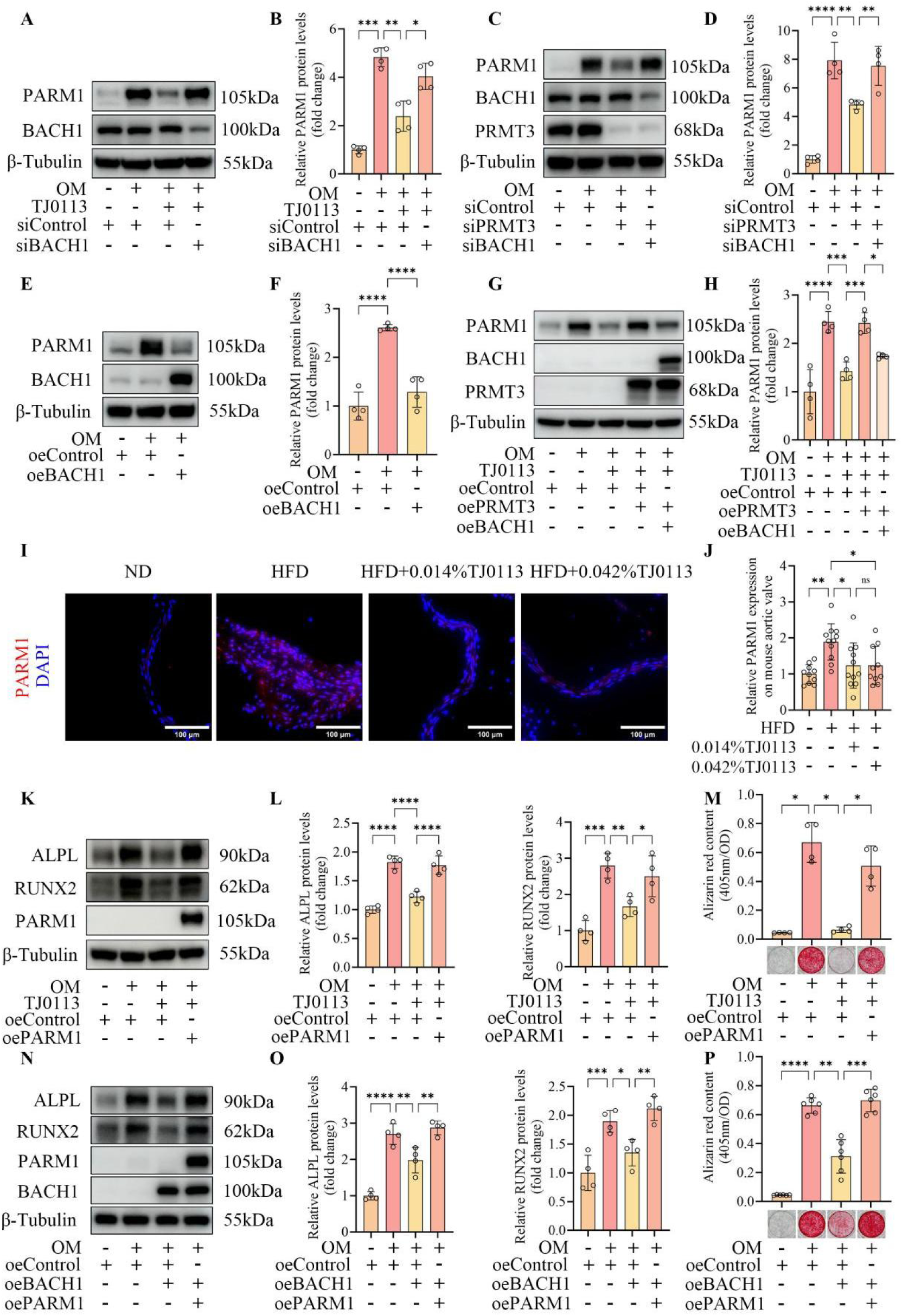
PARM1 acts as a downstream pro-osteogenic effector in CAVD. (A) Representative Western blot images of PARM1 and BACH1 in hVICs transfected with siControl or siBACH1 and cultured under different conditions. (B) Quantification of PARM1 protein levels (n = 4). (C) Representative Western blot images of PARM1, BACH1, and PRMT3 in hVICs transfected with siControl, siPRMT3 + siControl (as a control for siBACH1), or siPRMT3 + siBACH1, and cultured under different conditions. (D) Quantification of PARM1 protein levels (n = 4). (E) Representative Western blot images of PARM1 and BACH1 in hVICs transfected with oeBACH1 or oeControl adenovirus and cultured under different conditions. (F) Quantification of PARM1 protein levels (n = 4). (G) Representative Western blot images of PARM1, BACH1, and PRMT3 in hVICs transfected with oeControl, oePRMT3 + oeControl (as a control for oeBACH1), or oePRMT3 + oeBACH1 adenovirus, and cultured under different conditions. (H) Quantification of PARM1 protein levels (n = 4). (I) Representative immunofluorescence images of PARM1 in aortic valve leaflets of *LDLr^−/−^* mice from four groups: normal chow diet (ND), high-fat diet (HFD), HFD + 0.014% TJ0113, and HFD + 0.042% TJ0113, after 24 weeks of feeding; nuclei were counterstained with DAPI. (J) Quantification of PARM1 fluorescence intensity in mouse aortic valves (n = 10-11). (K) Representative Western blot images of ALPL, RUNX2, and PARM1 in hVICs transfected with oeControl or oePARM1 lentivirus and cultured under different conditions. (L) Quantification of ALPL and RUNX2 protein levels (n = 4). (M) Alizarin Red staining and quantification of calcium deposition in hVICs transfected with oeControl or oePARM1 lentivirus and cultured under different conditions for 21 days (n = 4). (N) Representative Western blot images of ALPL, RUNX2, PARM1, and BACH1 in hVICs transfected with oeControl, oeBACH1 + oeControl (as a control for oePARM1), or oeBACH1 + oePARM1, and cultured under different conditions. (O) Quantification of ALPL and RUNX2 protein levels (n = 4). (P) Alizarin Red staining and quantification of calcium deposition in hVICs transfected with oeControl, oeBACH1 + oeControl (as a control for oePARM1), or oeBACH1 + oePARM1 lentivirus, and cultured under different conditions for 21 days (n = 6). Scale bars: (I) 100 μm. For the statistical analysis, *P < 0.05, **P < 0.01, ***P < 0.001, and ****P < 0.0001 indicate significant differences, and ns indicates no significant difference.

## Discussion

To our knowledge, this is the first evidence that mitophagy is impaired in human calcified aortic valve tissue and during osteogenic differentiation of hVICs, as demonstrated by single-cell RNA sequencing, TEM, western blot, and the well-established mitophagy reporter mt-Keima. We establish that mitophagy deficiency is a critical driver of CAVD progression and, more importantly, demonstrate that TJ0113—an orally available mitophagy-activating compound with translational potential—attenuates disease progression both in vivo and in vitro. Mechanistically, PRMT3 is released from the mitophagy receptor MCL1 during hVIC osteogenic differentiation, leading to PRMT3 accumulation in the cytosol. This enhances PRMT3-BACH1 interaction, which in turn reduces BACH1 nuclear translocation. Consequently, BACH1-mediated transcriptional repression of *PARM1*, a key pro-osteogenic factor, is diminished, resulting in increased PARM1 expression and ultimately promoting hVIC osteogenic differentiation. These events constitute a dysregulated mitochondrial-to-nuclear communication initiated by impaired mitophagy and driving disease progression. Overall, this study establishes impaired mitophagy as a driver of CAVD, uncovers a novel mechanism, and positions mitophagy activation as a potential strategy to delay disease.

In the past decade, the importance of maintaining normal mitochondrial function in chronic diseases has received increasing attention. Mitochondrial dysfunction is now recognized as a hallmark of aging^33^. Similarly, in the context of CAVD, our previous work and emerging evidence have shown that mitochondrial dysfunction contributes to disease progression^6-10^. However, the role of mitophagy—a critical mitochondrial quality control process—remains poorly understood in CAVD. Although one study reported differential expression of some mitophagy-related proteins in the serum of CAVD patients, circulating protein levels are likely to differ substantially from those in aortic valve tissue, which is more directly relevant to the pathogenesis of aortic valve calcification^13^. A recent study implicated the mitophagy receptor BNIP3 as a protective factor during hVIC osteogenic differentiation in vitro, but did not examine how mitophagy is altered during this process, as appropriate controls were not included^14^. How mitophagy flux changes in calcified aortic valves and during hVIC osteogenic differentiation, the key cellular process driving CAVD, remains unclear. More importantly, whether modulating mitophagy affects CAVD progression has not been addressed in vivo, highlighting a critical gap in current understanding of CAVD pathogenesis. Therefore, prompted by the enrichment of the mitophagy pathway in our scRNA-seq data, we systematically evaluated the role of mitophagy in CAVD to address these gaps. Given the lack of effective pharmacologic therapies for CAVD in clinical practice, we selected TJ0113, a highly selective mitophagy-activating compound from our previous screening, for this study. This orally available agent has established safety in rats (26-week) and beagle dogs (39-week) toxicology studies^21^, and has reported positive phase II results in early-stage Parkinson’s disease (NCT06596005) and advanced to a phase III trial (NCT07741045), highlighting its translational potential. As a result, we revealed that mitophagy is impaired in human calcified aortic valves and during hVIC osteogenic differentiation, and that TJ0113 effectively activates MCL1-mediated mitophagy, restores mitochondrial structure and function, and attenuates CAVD progression.

MCL1, which is mainly localized to the mitochondrial outer membrane and has also been found in the mitochondrial matrix, has been characterized as a binding protein that sequesters and inactivates its partners before its identification as a mitophagy receptor^26, 27, 34^. Accordingly, after confirming that MCL1 is essential for the protective effect of TJ0113, we performed Co-IP followed by LC-MS/MS and identified PRMT3 as a potential downstream substrate. Traditionally, PRMT3 has been thought to localize mainly to the cytosol. However, a recent study reported its mitochondrial localization in Hepa 1-6, PLC-8024, and HEK293 cell lines^28^. Of note, our subcellular fractionation experiments revealed that PRMT3 is also present in the mitochondria of hVICs. More importantly, we observed that PRMT3 is released from MCL1 into the cytosol during hVIC osteogenic differentiation. These findings extend the substrate spectrum of MCL1 and provide new insights into mitochondrial-cytosolic protein communication.

PRMT3, a member of the protein arginine methyltransferase family, has been implicated in promoting osteogenic differentiation in mesenchymal stem cells and vascular smooth muscle cells^35, 36^. More recently, Xi Zhang et al. reported that in hVICs cultured with osteogenic medium supplemented with BMP-2, RUNX2 can recruit p300 to enhance H3K27ac enrichment at the PRMT3 promoter, driving PRMT3 transcription and protein upregulation. PRMT3 in turn dimethylates PCSK9 at arginine 582, protecting it from ubiquitin-mediated degradation and thereby promoting hVIC osteogenic differentiation^37^. They also found that PRMT3 knockdown in a CAVD mouse model alleviated disease progression, identifying PRMT3 as a risk factor in CAVD^37^. In our study, however, PRMT3 total protein levels remained unchanged during hVIC osteogenic differentiation. This discrepancy may be attributable to the fact that Xi Zhang et al. supplemented their osteogenic culture medium with BMP-2, one of the most potent osteogenic signals, whose upregulation occurs primarily at mid-to-late stages of differentiation^38, 39^. In addition, BMP-2, p300, and RUNX2 are known to form a complex regulatory network: BMP-2 enhances p300-mediated acetylation of RUNX2, increasing RUNX2 protein levels and transcriptional activity^40^; BMP-2 also upregulates p300 expression and enzymatic activity, while simultaneously increasing RUNX2 protein and enhancing RUNX2-p300 interaction^41^; conversely, RUNX2 recruits p300 to the BMP-2 promoter, promoting BMP-2 transcription^42^. These observations suggest a positive feedback loop among BMP-2, p300, and RUNX2 that synergistically amplifies osteogenic signaling. Compared with the study by Xi Zhang et al., our differentiation medium did not contain BMP-2 and reflected a relatively early stage of osteogenic differentiation. Nevertheless, PRMT3 remained a critical driver in our model, suggesting that PRMT3 contributes to hVIC osteogenic differentiation across multiple stages and through distinct mechanisms. Notably, our study further reveals that the pro-osteogenic function of PRMT3 is governed not only by its expression level but also by its subcellular distribution and protein-protein interaction status.

In probing PRMT3 downstream mechanism, the transcriptional repressor BACH1 emerged as a PRMT3-interacting protein. Indeed, other type I PRMT family members including PRMT4 and PRMT1 have been reported to directly interact with or regulate BACH1 in previous studies, supporting the plausibility of the PRMT3-BACH1 interaction^43, 44^. Importantly, our CUT&Tag data revealed a widespread loss of BACH1 genomic occupancy during osteogenic differentiation for the first time. This loss was attributed to enhanced PRMT3-BACH1 interaction, which restricted BACH1 nuclear translocation and was reversed by TJ0113-mediated mitophagy activation. Functionally, BACH1 has been shown to suppress osteogenic differentiation in two other cell types: its knockdown upregulated RUNX2 and BMP-6 in periodontal ligament cells^31^, as well as ALPL, RUNX2, and osteocalcin in bone marrow mesenchymal stem cells^32^, suggesting that BACH1 may function as a negative regulator of osteogenic differentiation. In our study, we confirmed for the first time that BACH1 serves as a protective transcription factor to inhibit hVIC osteogenic differentiation. To further investigate how BACH1 inhibits hVIC osteogenic differentiation, we identified PARM1 as a candidate downstream target through multi-omics integration of bulk RNA-seq, scRNA-seq, and CUT&Tag data. PARM1 was previously identified by our group as a key pro-osteogenic factor upregulated in calcified aortic valves and during hVIC osteogenic differentiation^22^. The findings in this study further extend our previous work by revealing why PARM1 significantly increases during hVIC osteogenic differentiation—the loss of BACH1 binding to the PARM1 promoter and the consequent relief of PARM1 transcriptional repression. Each of these interventions—mitophagy activation, PRMT3 knockdown, and BACH1 overexpression—can reduce PARM1 expression. Together with the rescue experiments which confirm PARM1 as the downstream effector of this axis, these findings establish a molecular link between mitophagy impairment and the expression of a CAVD-relevant gene.

Most studies on CAVD pathogenesis have focused on changes in protein expression levels. Beyond expression levels, our findings reveal that altered protein-protein interactions and subcellular localization play a critical role in promoting CAVD progression. Here, we identify a signaling cascade that constitutes a form of mitochondrial-to-nuclear communication: mitochondrial sequestration of an intermediate protein (PRMT3) controls its cytosolic availability, thereby regulating the nuclear translocation of transcription factor (BACH1) and its transcriptional repression of the pro-osteogenic factor (PARM1). This extends the current understanding of mitochondrial-to-nuclear signaling and highlights the importance of examining protein localization and interaction dynamics when investigating CAVD mechanisms.

Several limitations of this study should be acknowledged. First, although our in vitro and in vivo models mimic key features of CAVD, they do not fully reproduce the complex pathological process in humans. Second, we did not detect arginine methylation of BACH1 by PRMT3 in our IP experiments and LC-MS/MS analyses. However, this is consistent with a previous report showing that PRMT1, another type I arginine methyltransferase, also fails to methylate BACH1^44^. How PRMT3 regulates BACH1 nuclear translocation remains to be determined. Third, while our study establishes mitophagy impairment as a critical driver of CAVD, the upstream causes of mitophagy dysfunction remain to be fully elucidated. Further investigation into these mechanisms will deepen our understanding of CAVD pathogenesis.

In summary, this study identifies mitophagy impairment in hVICs as a critical driver of CAVD. For the first time, we reveal how mitophagy impairment transcriptionally regulates pro-osteogenic gene expression through the MCL1-PRMT3-BACH1-PARM1 axis, a dysregulated mitochondrial-to-nuclear communication. During osteogenic differentiation, MCL1-mediated sequestration of PRMT3 is weakened, leading to enhanced PRMT3-BACH1 interaction and subsequent inhibition of BACH1 nuclear translocation. This in turn diminishes BACH1-mediated transcriptional repression of pro-osteogenic PARM1, increasing PARM1 expression and promoting hVIC osteogenic differentiation. Importantly, we demonstrate that TJ0113, an orally available mitophagy activator with translational potential, activates mitophagy and thereby corrects this communication, attenuating disease progression. These findings establish selective mitophagy activation as a promising therapeutic strategy for CAVD.

## References

1. Desai MY, Braunwald E. The Pathophysiologic Basis and Management of Calcific Aortic Valve Stenosis: JACC State-of-the-Art Review. J AM COLL CARDIOL 2025; 86(9):659–672.

2. Small AM, Yutzey KE, Binstadt BA, Voigts Key K, Bouatia-Naji N, Milan D, et al. Unraveling the Mechanisms of Valvular Heart Disease to Identify Medical Therapy Targets: A Scientific Statement From the American Heart Association. CIRCULATION 2024; 150(6):e109–e128.

3. Xie M, Qian X, Yan G, Chen L, Ma R, Xiao Y. Global, regional, and national burden of non-rheumatic calcific aortic valve disease, 1990-2021: a systematic analysis for the Global Burden of Disease Study 2021. INT J SURG 2026; 112(2):3238–3251.

4. Otto CM, Newby DE, Hillis GS. Calcific Aortic Stenosis: A Review. JAMA-J AM MED ASSOC 2024; 332(23):2014–2026.

5. Qian X, Xu L, Geng B, Li F, Dong N. Navigating the Landscape of Translational Medicine of Calcific Aortic Valve Disease: Bridging Bench to Bedside. JACC Asia 2025; 5(4):503–515.

6. Chen J, Ren T, Xie L, Hu H, Li X, Maitusong M, et al. Enhancing aortic valve drug delivery with PAR2-targeting magnetic nano-cargoes for calcification alleviation. NAT COMMUN 2024; 15(1):557.

7. Liu F, Chen J, Hu W, Gao C, Zeng Z, Cheng S, et al. PTP1B Inhibition Improves Mitochondrial Dynamics to Alleviate Calcific Aortic Valve Disease Via Regulating OPA1 Homeostasis. JACC-BASIC TRANSL SC 2022; 7(7):697–712.

8. Liu H, Yin H, Wang Z, Yuan Q, Xu F, Chen Y, et al. Rho A/ROCK1 signaling-mediated metabolic reprogramming of valvular interstitial cells toward Warburg effect accelerates aortic valve calcification via AMPK/RUNX2 axis. CELL DEATH DIS 2023; 14(2):108.

9. Hu D, Lin Y, Huang H, Xian G, Chen Y, Li J, et al. CircZBTB44-Encoded Peptide ZBTB44-342aa Alleviates Aortic Valve Calcification Via cGAS-STING Inhibition. CIRC RES 2026; 138(3):e326771.

10. Song N, Ji E, Yu JE, Choi KH, Kim DH, Song JM, et al. Spermidine Enhances Mitochondrial Function and Mitigates Aortic Valve Calcification: Implications for DNA Methyltransferase-1 Activity. JACC-BASIC TRANSL SC 2025; 10(3):345–366.

11. Picca A, Faitg J, Auwerx J, Ferrucci L, D’Amico D. Mitophagy in human health, ageing and disease. NAT METAB 2023; 5(12):2047–2061.

12. Wang S, Long H, Hou L, Feng B, Ma Z, Wu Y, et al. The mitophagy pathway and its implications in human diseases. SIGNAL TRANSDUCT TAR 2023; 8(1):304.

13. Morciano G, Patergnani S, Pedriali G, Cimaglia P, Mikus E, Calvi S, et al. Impairment of mitophagy and autophagy accompanies calcific aortic valve stenosis favouring cell death and the severity of disease. CARDIOVASC RES 2022; 118(11):2548–2559.

14. Bian JH, Gu JX, Yue Z, Yuan CZ, Ji HB, Li A, et al. KLF5 Downregulation Links Impaired BNIP3-Mediated Mitophagy to Inflammatory Valve Remodeling in Calcific Aortic Valve Disease. ADV SCI 2026:e77368.

15. Cen X, Chen Y, Xu X, Wu R, He F, Zhao Q, et al. Pharmacological targeting of MCL-1 promotes mitophagy and improves disease pathologies in an Alzheimer’s disease mouse model. NAT COMMUN 2020; 11(1):5731.

16. Cen X, Xu X, Xia H. Targeting MCL1 to induce mitophagy is a potential therapeutic strategy for Alzheimer disease. AUTOPHAGY 2021; 17(3):818–819.

17. Wang K, Wu X, Li H, Xu X, Cheng F, Chen J, et al. Mitophagy Reprograms Lactate Metabolism to Suppress THBS1 via H3K18la Reduction, Alleviating Intervertebral Disc Degeneration. RESEARCH-CHINA 2025; 8:957.

18. Chen X, Xing S, Cen X, Shen J, Zhu J, Lin X, et al. Novel mitophagy activator TJ0113 restricts cytosolic mtDNA to suppress ZBP1-driven inflammation in acute lung injury. REDOX BIOL 2026; 96:104337.

19. Liu Z, Fang D, Chen K, Dong L, Huang H, Chen Z. Novel mitophagy inducer TJ0113 alleviates pulmonary inflammation during acute lung injury. FRONT PHARMACOL 2025; 16:1590458.

20. Huang C, Liao J, Cen X, Jiao C, Chen S, Liu D, et al. TJ0113-induced mitophagy in acute liver failure detected by Raman microspectroscopy. REDOX BIOL 2025; 83:103654.

21. Tian Y, Wang F, Shen J, Zhang R, Chen Q, Xu L, et al. Preclinical safety and toxicokinetic evaluation of TJ0113, a first-in-class mitophagy inducer for Parkinson’s disease. TOXICOL APPL PHARM 2025; 505:117565.

22. Hu H, Zhu X, Chen J, Cao N, Yang S, Xie L, et al. Knockdown of PARM1 Alleviates Aortic Valve Calcification via the PRKCH-MAPK Signaling Pathway. JACC-BASIC TRANSL SC 2025; 10(8):101260.

23. Iqbal F, Schlotter F, Becker-Greene D, Lupieri A, Goettsch C, Hutcheson JD, et al. Sortilin enhances fibrosis and calcification in aortic valve disease by inducing interstitial cell heterogeneity. EUR HEART J 2023; 44(10):885–898.

24. Antico O, Thompson PW, Hertz NT, Muqit M, Parton LE. Targeting mitophagy in neurodegenerative diseases. NAT REV DRUG DISCOV 2025; 24(4):276–299.

25. Wang Q, Sun Y, Li TY, Auwerx J. Mitophagy in the pathogenesis and management of disease. CELL RES 2026; 36(1):11–37.

26. Widden H, Placzek WJ. The multiple mechanisms of MCL1 in the regulation of cell fate. COMMUN BIOL 2021; 4(1):1029.

27. Sancho M, Leiva D, Lucendo E, Orzáez M. Understanding MCL1: from cellular function and regulation to pharmacological inhibition. FEBS J 2022; 289(20):6209–6234.

28. Shi Y, Wu Z, Liu S, Zuo D, Niu Y, Qiu Y, et al. Targeting PRMT3 impairs methylation and oligomerization of HSP60 to boost anti-tumor immunity by activating cGAS/STING signaling. NAT COMMUN 2024; 15(1):7930.

29. Wang Y, Wang C, Guan X, Ma Y, Zhang S, Li F, et al. PRMT3-Mediated Arginine Methylation of METTL14 Promotes Malignant Progression and Treatment Resistance in Endometrial Carcinoma. ADV SCI 2023; 10(36):e2303812.

30. Shi Y, Niu Y, Yuan Y, Li K, Zhong C, Qiu Z, et al. PRMT3-mediated arginine methylation of IGF2BP1 promotes oxaliplatin resistance in liver cancer. NAT COMMUN 2023; 14(1):1932.

31. Yuan Z, Li J, Zou X, Liu C, Lu J, Ni C, et al. Knockdown of Bach1 protects periodontal bone regeneration from inflammatory damage. J CELL MOL MED 2023; 27(22):3465–3477.

32. Xiao S, Li G, Tan M, Liu W, Li W. Loss of BACH1 improves osteogenic differentiation in glucocorticoid-induced hBMSCs through restoring autophagy. BMC MUSCULOSKEL DIS 2024; 25(1):665.

33. Kroemer G, Maier AB, Cuervo AM, Gladyshev VN, Ferrucci L, Gorbunova V, et al. From geroscience to precision geromedicine: Understanding and managing aging. CELL 2025; 188(8):2043–2062.

34. Escudero S, Zaganjor E, Lee S, Mill CP, Morgan AM, Crawford EB, et al. Dynamic Regulation of Long-Chain Fatty Acid Oxidation by a Noncanonical Interaction between the MCL-1 BH3 Helix and VLCAD. MOL CELL 2018; 69(5):729–743.

35. Min Z, Xiaomeng L, Zheng L, Yangge D, Xuejiao L, Longwei L, et al. Asymmetrical methyltransferase PRMT3 regulates human mesenchymal stem cell osteogenesis via miR-3648. CELL DEATH DIS 2019; 10(8):581.

36. Zhou G, Zhang C, Peng H, Su X, Huang Q, Zhao Z, et al. PRMT3 methylates HIF-1α to enhance the vascular calcification induced by chronic kidney disease. MOL MED 2024; 30(1):8.

37. Zhang X, Hao Y, Han D, Jin X, Shang X, Zhang L, et al. PRMT3-Mediated Arginine Methylation Stabilizes PCSK9 to Promote Aortic Valve Calcification. CIRCULATION 2026.

38. Ryoo HM, Lee MH, Kim YJ. Critical molecular switches involved in BMP-2-induced osteogenic differentiation of mesenchymal cells. GENE 2006; 366(1):51–57.

39. Migliorini E, Valat A, Picart C, Cavalcanti-Adam EA. Tuning cellular responses to BMP-2 with material surfaces. CYTOKINE GROWTH F R 2016; 27:43–54.

40. Jeon EJ, Lee KY, Choi NS, Lee MH, Kim HN, Jin YH, et al. Bone morphogenetic protein-2 stimulates Runx2 acetylation. J BIOL CHEM 2006; 281(24):16502–16511.

41. Jun JH, Yoon WJ, Seo SB, Woo KM, Kim GS, Ryoo HM, et al. BMP2-activated Erk/MAP kinase stabilizes Runx2 by increasing p300 levels and histone acetyltransferase activity. J BIOL CHEM 2010; 285(47):36410–36419.

42. Tsai SY, Huang YL, Yang WH, Tang CH. Hepatocyte growth factor-induced BMP-2 expression is mediated by c-Met receptor, FAK, JNK, Runx2, and p300 pathways in human osteoblasts. INT IMMUNOPHARMACOL 2012; 13(2):156–162.

43. He Y, Guo J, Yu Y, Jin J, Jiang Q, Li Q, et al. BACH1 regulates the differentiation of vascular smooth muscle cells from human embryonic stem cells via CARM1-mediated methylation of H3R17. CELL REP 2023; 42(12):113468.

44. Nguyen C, Colón-Emeric BA, Murakami S, Shujath M, Yi C. PRMT1 promotes epigenetic reprogramming associated with acquired chemoresistance in pancreatic cancer. CELL REP 2024; 43(5):114176.

