## Supplementary materials for "Pharmacological activation of mitophagy prevents calcific aortic valve disease by targeting dysregulated mitochondrial-to-nuclear communication"

Hongguang Xia, PhD.

Research Center of Clinical Pharmacy of The First Affiliated Hospital & Liangzhu Laboratory, Zhejiang University School of Medicine, Hangzhou, 311121, China

Jinghai Chen, PhD.

Department of Cardiology, The Second Affiliated Hospital, School of Medicine, Zhejiang University, Hangzhou 310009, China

[†] Contributed equally to the manuscript.

#### Supplementary materials

#### Cell culture and treatment

Primary human valvular interstitial cells (hVICs) were isolated from non-calcified aortic valves. Briefly, aortic valve leaflets were washed four times with phosphate-buffered saline (PBS), digested with high-glucose DMEM supplemented with 2 mg/mL collagenase I (Sigma, SCR103) for 15 min, and then vortexed to remove valvular endothelial cells. The valve leaflets were then cut into 1-2 mm pieces and digested with 1 mg/mL collagenase I at 37°C in a 5% $CO_2$ incubator for 12 hours. The tissue fragments were further disrupted by repeated pipetting, and digestion was stopped by adding an adequate volume of complete growth medium. The suspension was centrifuged at 600× g for 10 minutes. Finally, isolated hVICs were resuspended and cultured in high-glucose DMEM supplemented with 10% fetal bovine serum (FBS) and 1% penicillin/streptomycin (complete growth medium, CGM). After expansion, cells from the third to seventh passage were used for experiments. For osteogenic induction, hVICs were cultured in osteogenic medium (OM; CGM supplemented with 10 mM β-glycerophosphate, 0.25 mM ascorbic acid, and 10 nM dexamethasone) for 4 days[1, 2]. For pharmacological activation of mitophagy, hVICs were treated with 5 μM TJ0113 under OM culture conditions. The respective media were changed every day.

#### Single-cell RNA sequencing

Calcified and non-calcified human aortic valve leaflets (one leaflet per sample, from two patients per group) were washed four times in ice-cold PBS, minced into approximately 1 mm pieces, and digested in high-glucose DMEM supplemented with 1 mg/mL collagenase I (Sigma, SCR103) and 1% penicillin/streptomycin for 12 hours at 37°C in a 5% $CO_2$ incubator. The digested tissue was further dissociated by repeated pipetting, and digestion was terminated by adding an adequate volume of CGM. The cell suspension was centrifuged at 600× g for 5 minutes, and the pellet was resuspended in CGM. Cell suspensions were transported on ice to LC-BIO Technology (Hangzhou, China), where subsequent procedures including filtration, red blood cell lysis, flow cytometric sorting of viable cells, and quality control (viability > 80%, aggregation rate < 20%) were performed. Single-cell capture and barcoding were carried out on the Chromium iX platform (10X Genomics). cDNA amplification and library construction were performed using the Chromium GEM-X Single Cell 3′ Kit (10X Genomics) according to the manufacturer's protocol. Libraries were quality-controlled using the Agilent dsDNA 915 Reagent Kit (Agilent, DNF-915-K0500). Sequencing was performed on the Illumina NovaSeq X Plus platform (PE150) with a minimum depth of 20,000 reads per cell.

Raw sequencing data were aligned and gene-expression count matrices were generated using Cell Ranger 8.0.1. Analysis was performed using the Seurat package (version 4.3.0.1). Cells were filtered based on the following criteria: UMI count < 10, number of genes per cell < 200, or mitochondrial gene percentage > 30%. Genes expressed in fewer than three cells were excluded. Integration of samples across conditions was performed using the Harmony method. Principal component analysis was conducted based on the 2,000 most variable genes to identify nearest neighbors, and clustering was performed using the FindCluster function. hVIC populations were identified based on marker gene lists generated by the FindAllMarkers function. Differentially expressed genes were identified using $|\log_2 FC| > 0.5$ and adjusted $P < 0.05$, followed by enrichment analysis.

For hVIC scRNA-seq, cells cultured in CGM, OM, or OM supplemented with TJ0113 for four days (n = 2 per group) were harvested using trypsin-EDTA (Cienry, Zhejiang, China). Single-cell suspensions were prepared and subjected to the same library preparation and sequencing procedures as described for the tissue samples.

**Transmission electron microscopy**

Transmission electron microscopy (TEM) was used to observe the mitochondrial ultrastructure and mitophagy. For TEM observation of aortic valves, calcific or non-calcified aortic valves were collected in cold PBS immediately after surgery. The tissue was fully rinsed, cut into 1-2 mm pieces using a sharp surgical blade, and immediately fixed in 2.5% glutaraldehyde (Solarbio, Beijing, China). After complete fixation, the samples were processed and observed at the Center of Cryo-Electron Microscopy (CCEM), Zhejiang University. For TEM observation of hVIC mitochondrial structure, hVICs cultured in CGM, OM, or OM supplemented with TJ0113 for four days were harvested using trypsin-EDTA. For TEM observation of hVIC mitophagy, bafilomycin A1 was added 6 hours before harvest. All images were obtained using a Thermo Scientific Talos L120C at 13,500× magnification.

**Transfection of siRNA**

hVICs were transfected with small interfering RNA (siRNA) using Lipofectamine™ RNAiMAX Transfection Reagent (Invitrogen, 13778150) according to the manufacturer's instructions. The siRNAs targeting *MCL1*, *PINK1*, *PRMT3*, *BACH1*, and negative control were purchased from GenePharma (Suzhou, China). The sequences of these siRNAs are listed in Supplementary Table 1.

**Adenovirus-mediated gene overexpression**

Adenovirus overexpressing human *PRMT3* (3×Flag-tagged) or *BACH1* (HA-tagged) was designed and purchased from GENTIMES BIO (Guangzhou, China). After reaching 60% confluence, hVICs were counted and incubated with adenovirus in half the culture volume of DMEM (containing 10% FBS) at a multiplicity of infection (MOI) of 40 for four hours. Then, another half volume of fresh DMEM containing 10% FBS was added to each well, and incubation was continued for 12 hours. After switching to fresh CGM for two days, hVICs were treated with different media according to the experimental groups.

**Mitophagy level evaluation with mt-Keima**

Mitophagy level in hVICs was evaluated using mt-Keima, a chimeric protein previously established as a robust sensor of mitophagy[3-5]. mt-Keima is a fusion protein consisting of the fluorescent protein Keima and a mitochondrial signal sequence derived from cytochrome c oxidase subunit IV, which allows its specific expression in mitochondria. The excitation wavelength of mt-Keima under neutral conditions is 438 nm. When mitochondria are exposed to the acidic environment of lysosomes (i.e., when mitophagy occurs), the excitation wavelength shifts to 550 nm. The adenovirus expressing mt-Keima used in this study was purchased from Hanbio (Shanghai, China).

hVICs were seeded at a density of $3 \times 10^5$ cells per well in 6-well plates. On the second day, hVICs were transfected with the mt-Keima adenovirus using a method similar to that described above (MOI of 5). After switching to fresh CGM for two days, hVICs were treated with different media according to the experimental groups. Finally, hVICs were harvested, and mt-Keima was evaluated by flow cytometry (BD Biosciences) as described in previous studies[6, 7].

**Lentivirus transfection**

Control lentivirus, gene-specific short hairpin RNA (shRNA) lentivirus, and gene overexpression lentivirus were used in hVICs calcium deposition experiments. After determining the optimal MOI for each lentivirus, hVICs were seeded at a density of $5 \times 10^4$ cells per well in 12-well plates and were transfected with lentivirus using HitransG P (Genechem, Shanghai, China) for 12 hours. After switching to fresh CGM for two days, hVICs were treated with different media according to the experimental groups. *PARM1* overexpression lentivirus was also used in rescue experiments to assess osteogenic differentiation markers in hVICs.

#### Mitochondrial membrane potential evaluation

The mitochondrial membrane potential of hVICs was evaluated using JC-1 (Beyotime, C2003S). Briefly, hVICs were incubated with JC-1 probe at 37°C for 30 min, washed three times, and then detached using trypsin-EDTA. The detached cells were resuspended in ice-cold HBSS and immediately analyzed by flow cytometry.

#### Reactive oxygen species (ROS) measurement

After three washes with PBS, hVICs were incubated with DMEM supplemented with 10 μM DCFH-DA (Sigma, D6883) at 37°C for 20 min. Then hVICs were washed three times, detached using trypsin-EDTA, resuspended in ice-cold HBSS, and immediately analyzed by flow cytometry.

#### Alkaline phosphatase activity assessment

Alkaline phosphatase (ALP) activity of hVICs was assessed using an ALP activity assay kit (Beyotime, P0321). Briefly, hVICs were lysed using lysis buffer without phosphatase inhibitors (Beyotime, P0013J). The cell lysates were then incubated with p-nitrophenyl phosphate buffer solution in a 96-well plate. After incubation at 37°C for 10 min, stop solution was added to each well, and the absorbance at 405 nm was measured. ALP activity was calculated based on a standard curve generated from serial dilutions of p-nitrophenol and normalized to total protein concentration measured by a BCA protein assay kit (Thermo Fisher, 23227).

**Alizarin red staining**

Alizarin red staining was performed to evaluate calcium deposition in hVICs after 21 days of different treatments. The respective media were changed every 2-3 days. On the day of staining, hVICs were washed with PBS and then fixed in 4% paraformaldehyde for 30 minutes. After washing three times with PBS, the cells were stained with alizarin red solution (Oricell, ALIR-10001) at room temperature for 20 minutes. Following three additional washes, calcification was photographed and visualized. For quantification, calcium nodules stained with alizarin red were dissolved in 10% acetic acid for 30 minutes, and the absorbance of the supernatant was measured at 405 nm.

**Dual-luciferase reporter assay**

The pRL-TK plasmid, PGL3-basic plasmid, human *PARM1* promoter plasmid (2000bp, in PGL3 vector), *BACH1* overexpression plasmid (oeBACH1), and the corresponding empty vector plasmid (oeControl) were constructed and validated by Tsingke (Beijing, China). The activity of the human PARM1 promoter plasmid was first confirmed in HEK293T cells in preliminary experiments. Subsequently, the luciferase reporter plasmids were co-transfected into HEK293T cells together with either the BACH1 overexpression plasmid or the empty vector. Forty-eight hours after transfection, cells were harvested using the passive lysis buffer provided in the kit, and luminescence was measured using a Spark multimode microplate reader (Tecan,

Männedorf, Switzerland) according to the protocol of the Dual-Luciferase Reporter Assay System (Promega, E1910).

**Quantitative real-time polymerase chain reaction (RT-qPCR)**

RNA of hVICs was extracted using EZ-press RNA Purification Kit (EZBioscience, B0004DP) following the manufacturer's protocol. The concentration and quality of RNA samples were evaluated using NanoDrop 2000. Subsequently, complementary DNA (cDNA) was synthesized using the PrimeScript RT kit (Takara, RR036). Then, RT-qPCR was performed on the LightCycler 480II (Roche, Switzerland) using TB Green Premix Ex Taq (cat #RR420A, Takara). Relative mRNA expression was calculated using the comparative Ct (ΔΔCt) method and normalized to *TUBB*. The sequences of primers were shown in the Supplementary Table 2.

**Western blot**

Total protein was extracted using RIPA buffer (Beyotime, P0013K) containing protease and phosphatase inhibitor cocktails (Fdbio Science, FD1001 and FD1002). Protein concentration was determined using a BCA protein assay kit (Thermo Fisher, 23227). Equal amounts of protein were separated by sodium dodecyl sulfate-polyacrylamide gel electrophoresis (SDS-PAGE) and transferred to 0.2 μm PVDF membranes (MilliporeSigma). After blocking with 5% bovine serum albumin (BSA) in Tris Buffered Saline with Tween-20 (TBST) at room temperature for 1 hour, the membranes were incubated with primary antibodies at 4°C overnight. Following three washes with TBST, the membranes were incubated with HRP-conjugated secondary antibodies (CST, 7074 or 7076) at room temperature for 1 hour and then washed three more times. The blots were visualized using an enhanced chemiluminescence reagent (Vazyme, E423) and an ImageQuant 800 system (Amersham, GE Healthcare). Band intensities were analyzed using ImageJ software. All primary antibodies used in this study are listed in Supplementary Table 3.

**Co-Immunoprecipitation (Co-IP)**

Cells were harvested using NP-40 lysis buffer (Beyotime, P0013F) containing protease and phosphatase inhibitor cocktails. The lysates were centrifuged at 12,000× g for 5 min at 4°C. The supernatant was collected, and an input sample was retained. The remaining supernatant was incubated with primary antibodies overnight at 4°C with rotation. The next day, protein A/G magnetic beads (MedChemExpress, HY-K0202) were washed with TBS four times. The antibody-protein solution was then incubated with the magnetic beads, and the mixture was rotated at room temperature for 1 hour. Afterward, the magnetic beads bound with antibody-protein immunocomplexes were washed four times with cold TBS containing protease and phosphatase inhibitor cocktails. Finally, the immunocomplexes were eluted by heat denaturation in SDS-PAGE loading buffer, and then analyzed by western blot.

**Co-IP followed by liquid chromatography-tandem mass spectrometry (LC-MS/MS)**

Co-IP followed by LC-MS/MS was performed to identify candidate proteins interacting with MCL1 or PRMT3. The Co-IP was carried out as described above, and the magnetic beads bound with protein-antibody immunocomplexes were rapidly frozen in liquid nitrogen. The bead samples were then transferred on dry ice to Shanghai Bioprofile Technology (Shanghai, China) for LC-MS/MS analysis. Given that the protein functions of MCL1 or PRMT3 depend on protein-protein interactions, candidate proteins were defined as those that were absent in the IgG control group and showed consistent detection patterns across three biological replicates within each experimental group (either all detected or all undetected), exhibited the same detection outcome in the CGM and TJ0113 groups but the opposite outcome in the OM group, and were validated by reliable MS/MS spectra.

**Molecular docking**

Molecular docking in this study was performed by Phadcalc (Chengdu, China). Briefly, protein-protein docking was carried out with the HDOCK program, and binding models were calculated based on the iterative scoring functions ITScorePP and ITScorePR. Referring to the docking score distribution of complexes resolved in the Protein Data Bank (PDB) (typically around -200 or lower), a confidence score was introduced based on the HDOCK program to evaluate binding probability. The confidence score was calculated using the following formula: Confidence score = $1.0 / [1.0 + e^{0.02 \times (\text{Docking\_Score} + 150)}]$. A score > 0.7 indicates a highly likely interaction, a score between 0.5 and 0.7 suggests a possible interaction, and a score < 0.5 indicates a low binding probability.

**Isolation of mitochondria and cytosol**

Mitochondria were isolated from hVICs using a mitochondrial isolation and protein extraction kit (Proteintech, PK10016). Soluble cytosolic proteins were extracted based on previous studies[8, 9]. Briefly, hVICs were harvested using trypsin-EDTA, washed with PBS, and incubated with digitonin solution containing 35 μg/mL digitonin, 150 mM NaCl, 50 mM HEPES, and protease and phosphatase inhibitor cocktails at 4°C with rotation for 10 min. The cell suspension was then centrifuged at 950× g for 5 min at 4°C. The supernatant was collected and further centrifuged at 17,000× g for 10 min at 4°C. The resulting supernatant, representing the cytosolic fraction, was collected for analysis.

**Isolation of nuclear and cytoplasmic fractions**

Nuclear and cytoplasmic fractionation was performed using a hypotonic lysis-based method[10-12]. Briefly, hVICs were harvested, washed with PBS, resuspended in hypotonic buffer (20 mM Tris-HCl [pH 7.4], 10 mM NaCl, 0.5 mM $MgCl_2$) supplemented with protease and phosphatase inhibitor cocktails. After incubation on ice for 10 min, DTT was added to a final concentration of 50 μM, and the mixture was incubated for an additional 10 min on ice. Subsequently, 10% NP-40 and 10% Tween-20 were added to final concentrations of 0.5% and 0.1%, respectively. The sample was vigorously vortexed for 10 s and then homogenized by 10 passages through a 27-gauge needle. A 1 μL aliquot of the homogenate was mixed with trypan blue, and microscopic examination confirmed that the nuclei had been released and remained intact. The homogenate was then centrifuged at 4,000 rpm for 10 min at 4°C, and the supernatant was collected as the cytoplasmic fraction. The nuclear pellet was washed twice with hypotonic buffer. For each wash, the pellet was resuspended in hypotonic buffer and centrifuged at 3,500 rpm for 5 min at 4°C, and the supernatant was carefully removed. The washed pellet was then lysed in ice-cold RIPA buffer supplemented with protease and phosphatase inhibitor cocktails. The lysate was vigorously vortexed every 3–4 min for a total of 30 min, followed by centrifugation at 16,000 × g for 10 min at 4°C. The resulting supernatant was collected as the nuclear fraction.

**Cleavage Under Targets and Tagmentation (CUT&Tag) sequencing**

We profiled the genome-wide DNA binding landscape of BACH1 in hVICs using CUT&Tag. hVICs from different groups were harvested by trypsinization, and all subsequent steps were performed by Jiayin Biotechnology Ltd. (Shanghai, China), which provided all reagents except the primary antibody. Briefly, $5 \times 10^5$ cells per reaction were bound to ConA magnetic beads (Bangs Laboratories) for 10 min at room temperature. The cell-bead complexes were then incubated overnight at 4°C with an anti-BACH1 primary antibody (CST, 33059), followed by a 60-min incubation with a goat anti-rabbit secondary antibody (N269; 1:100 dilution) and a 1-hour incubation with the pA-Tn5 fusion protein (1:100 dilution) at room temperature. Tagmentation was performed in a buffer containing 10 mM $MgCl_2$ for 1 hour at 37°C, and the reaction was terminated by adding SDS to a final concentration of 0.2% and incubating at 55°C for 10 min. Libraries were prepared using the NEBNext HiFi 2× PCR Master Mix and sequenced on the NovaSeq 6000 platform (PE150). Genome-wide binding signal analysis was performed using deepTools. To generate the heatmap, all genes were normalized by gene length. Sequencing signals were then calculated for each base pair across a region spanning from 3 kb upstream of the transcription start site (TSS) to 3 kb downstream of the transcription end site (TES). Differential peaks were identified using DEGseq ($|\log_2 FC| > 1$, $p < 0.05$).

**Bulk RNA sequencing**

Three days after transfection with siBACH1 or siControl, hVICs were lysed in TRIzol reagent (Invitrogen), and the lysates were rapidly frozen in liquid nitrogen and transferred on dry ice to LC-Bio Technology (Hangzhou, China) for RNA extraction and sequencing. After total RNA extraction, poly(A) enrichment was performed, followed by fragmentation, reverse transcription, and strand-specific cDNA library preparation. The libraries were sequenced on an Illumina NovaSeq 6000 platform (PE150). Raw reads were trimmed using fastp software and aligned to the GRCh38 genome with HISAT2. Transcript abundance was quantified as FPKM using StringTie. In differential expression analysis, genes with $|\log_2 FC| > 1$ and an adjusted p-value $<$ 0.05 were considered significantly differentially expressed.

#### Animal studies

##### High-fat diet-induced CAVD mouse model

Eight-week-old male *LDLr*$^{-/-}$ mice on a C57BL/6J background were fed a high-fat diet (TD-88137, SYSE Bio-tech, Changzhou, China) for 24 weeks to induce CAVD[13-15]. For pharmacological activation of mitophagy, TJ0113 was uniformly mixed into the high-fat diet by SYSE Bio-tech at two concentrations (0.014% and 0.042%), corresponding to low-dose and high-dose oral administration, respectively[16]. The feed was refreshed every two or three days. After 24 weeks, echocardiography was performed to evaluate aortic valve hemodynamics. Mice were then sacrificed after intraperitoneal injection of an overdose of sodium pentobarbital (0.2 g/kg). Plasma was collected for subsequent measurement of total cholesterol, low-density lipoprotein (LDL), triglycerides, blood glucose, creatinine, BUN, ALT, and AST. Heart samples were harvested for histological evaluation.

##### Aortic valve wire injury model

Aortic valve wire injury followed by a 12-week high-fat diet was performed in 8-week-old male *LDLr*$^{-/-}$ mice on a C57BL/6J background to induce CAVD. The procedure was performed as described previously[15, 17]. Briefly, a wire (ASAHI MIRACLEbros 6) was introduced into the right common carotid artery, advanced over the aortic valve, and inserted into left ventricle under echocardiographic guidance. Aortic valve wire injury was then induced by 20 scratches and 100 rotations of the wire. Three days after the procedure, mice were fed different diets according to their groups. Mice in the sham group underwent a sham procedure (right common carotid artery was exposed and ligated) and were fed a normal chow diet. Twelve weeks after the wire injury procedure, echocardiography was performed to evaluate aortic valve hemodynamics, and heart samples were harvested for histological evaluation.

**Echocardiography**

Transthoracic echocardiography was performed using a VisualSonics Vevo 3100 ultrasound system. The mice were maintained under light sedation with 0.5-1.5% isoflurane anesthesia, and heart rate was kept at 450-600 beats/min. Pulsed-wave Doppler was used to assess aortic valve hemodynamics, and M-mode imaging was used to evaluate cardiac function[14]. Each measurement was averaged over three consecutive cardiac cycles. All ultrasound data were acquired by the same animal sonographer.

**Histological analysis**

After transcardial perfusion and rinsing with PBS, mouse hearts were immersed in 30% sucrose solution for 24 hours and then embedded in OCT compound (Sakura Finetek). To ensure that all three aortic valve leaflets were visible in the same section, slices were cut perpendicular to the aortic root[15]. Hematoxylin and eosin (H&E) staining was performed to quantify aortic valve leaflet thickness, as described previously[15]. Alizarin red staining was performed using Alizarin Red S Solution

(Solarbio, G1450).

**Immunofluorescence staining**

Frozen sections were fixed in 4% paraformaldehyde for 10 min, permeabilized with 0.2% Triton X-100 for 20 min, blocked with 5% BSA (containing 0.1% Triton X-100) for 60 min, and incubated with specific primary antibodies overnight at 4°C. After four washes with PBS, sections were incubated with secondary antibodies (Invitrogen, A-31572, 1:400) at room temperature for 60 minutes. Following four additional washes with PBS, nuclei were counterstained with a mounting medium containing DAPI (Vector Laboratories, H-1200). Images were acquired using a Leica fluorescence microscope and analyzed with ImageJ software.

**Statistical Analysis**

Statistical analysis was performed using GraphPad Prism version 10. All quantitative experiments were performed with at least three biological replicates. Data are presented as mean ± standard deviation. Normality was assessed using the Shapiro-Wilk test. For comparisons between two groups, the unpaired Student's t-test was used for normally distributed data, and the Mann-Whitney U test was used for non-normally distributed data. For comparisons involving three or more groups, one-way ANOVA followed by Tukey's post hoc test was applied when data were normally distributed with homogeneous variances. When variances were unequal, the Brown-Forsythe test was used followed by Dunnett's T3 post hoc test. For non-normally distributed data across multiple groups, the Kruskal-Wallis test was performed with Dunn's post hoc test. A two-tailed p-value $< 0.05$ was considered statistically significant.

### References:


1. Rogers MA, Bartoli-Leonard F, Zheng KH, Small AM, Chen HY, Clift CL, et al. Major Facilitator Superfamily Domain Containing 5 Inhibition Reduces Lipoprotein(a) Uptake and Calcification in Valvular Heart Disease. *CIRCULATION* 2024; 149(5):391-401.
2. Liu F, Chen J, Hu W, Gao C, Zeng Z, Cheng S, et al. PTP1B Inhibition Improves Mitochondrial Dynamics to Alleviate Calcific Aortic Valve Disease Via Regulating OPA1 Homeostasis. *JACC-BASIC TRANSL SC* 2022; 7(7):697-712.
3. Antico O, Thompson PW, Hertz NT, Muqit M, Parton LE. Targeting mitophagy in neurodegenerative diseases. *NAT REV DRUG DISCOV* 2025; 24(4):276-299.
4. Wang Q, Sun Y, Li TY, Auwerx J. Mitophagy in the pathogenesis and management of disease. *CELL RES* 2026; 36(1):11-37.
5. Wang S, Long H, Hou L, Feng B, Ma Z, Wu Y, et al. The mitophagy pathway and its implications in human diseases. *SIGNAL TRANSDUCT TAR* 2023; 8(1):304.
6. Gilsrud AJ, Narendra DP. Methods for Monitoring Mitophagy Using mt-Keima. *Methods Mol Biol* 2024; 2845:151-160.
7. Winsor NJ, Killackey SA, Philpott DJ, Girardin SE. An optimized procedure for quantitative analysis of mitophagy with the mtKeima system using flow cytometry. *BIOTECHNIQUES* 2020; 69(4):249-256.
8. Holden P, Horton WA. Crude subcellular fractionation of cultured mammalian cell lines. *BMC RES NOTES* 2009; 2:243.
9. Nguyen M, Collier JJ, Ignatenko O, Morin G, Goyon V, Janer A, et al. MAPL regulates gasdermin-mediated release of mtDNA from lysosomes to drive pyroptotic cell death. *NAT CELL BIOL* 2025; 27(10):1708-1724.
10. DeCaprio J, Kohl TO. Using Dounce Homogenization to Lyse Cells for Immunoprecipitation. *Cold Spring Harb Protoc* 2019; 2019(7).
11. Huynh HT, Shcherbinina E, Huang HC, Rezaei R, Sarshad AA. Biochemical Separation of Cytoplasmic and Nuclear Fraction for Downstream Molecular Analysis. *Curr Protoc* 2024; 4(5):e1042.

12. Senichkin VV, Prokhorova EA, Zhivotovsky B, Kopeina GS. Simple and Efficient Protocol for Subcellular Fractionation of Normal and Apoptotic Cells. *CELLS-BASEL* 2021; 10(4).

13. Wang Y, Xu F, Song C, Wang X, Zhou H, Qu T, et al. Cyclic guanosine monophosphate-protein kinase G signaling attenuates aortic valve calcification through ULK1-mediated autophagy. *SIGNAL TRANSDUCT TAR* 2026; 11(1).

14. Chen J, Ren T, Xie L, Hu H, Li X, Maitusong M, et al. Enhancing aortic valve drug delivery with PAR2-targeting magnetic nano-cargoes for calcification alleviation. *NAT COMMUN* 2024; 15(1):557.

15. Hu H, Zhu X, Chen J, Cao N, Yang S, Xie L, et al. Knockdown of PARM1 Alleviates Aortic Valve Calcification via the PRKCH-MAPK Signaling Pathway. *JACC-BASIC TRANSL SC* 2025; 10(8):101260.

16. Tian Y, Wang F, Shen J, Zhang R, Chen Q, Xu L, et al. Preclinical safety and toxicokinetic evaluation of TJ0113, a first-in-class mitophagy inducer for Parkinson's disease. *TOXICOL APPL PHARM* 2025; 505:117565.

17. Iqbal F, Schlotter F, Becker-Greene D, Lupieri A, Goettsch C, Hutcheson JD, et al. Sortilin enhances fibrosis and calcification in aortic valve disease by inducing interstitial cell heterogeneity. *EUR HEART J* 2023; 44(10):885-898.

**Supplementary Figure 1. Single-cell RNA-sequencing analysis of the hVIC population from human calcified and non-calcified aortic valves.**

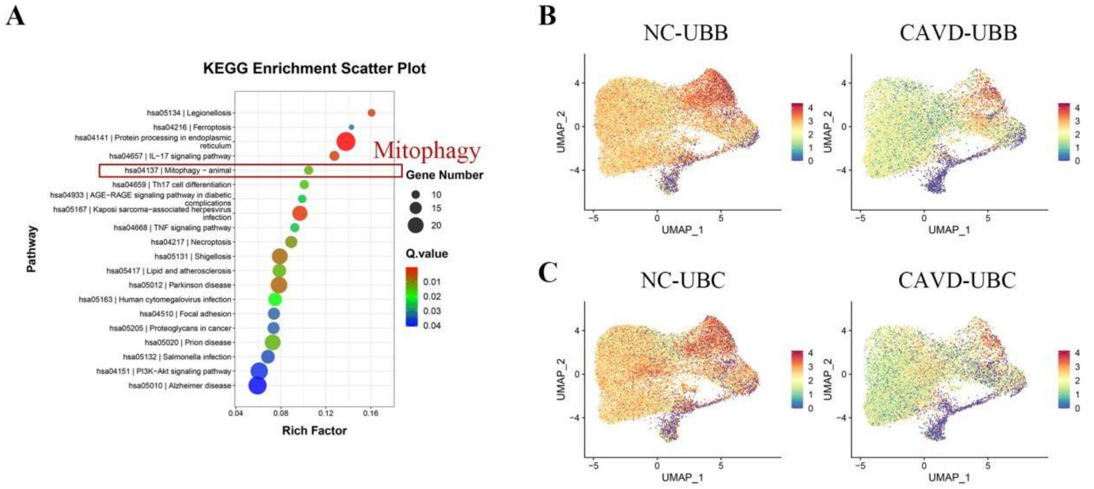


(A) KEGG pathway enrichment analysis of all differentially expressed genes in hVICs from calcified aortic valves, showing significant enrichment of the mitophagy pathway. (B-C) UMAP plots showing the downregulation of UBB (B) and UBC (C) in hVICs from calcified aortic valves compared with non-calcified valves.

**Supplementary Figure 2. Dose screening, safety evaluation, and effectiveness assessment of TJ0113 in hVICs.**

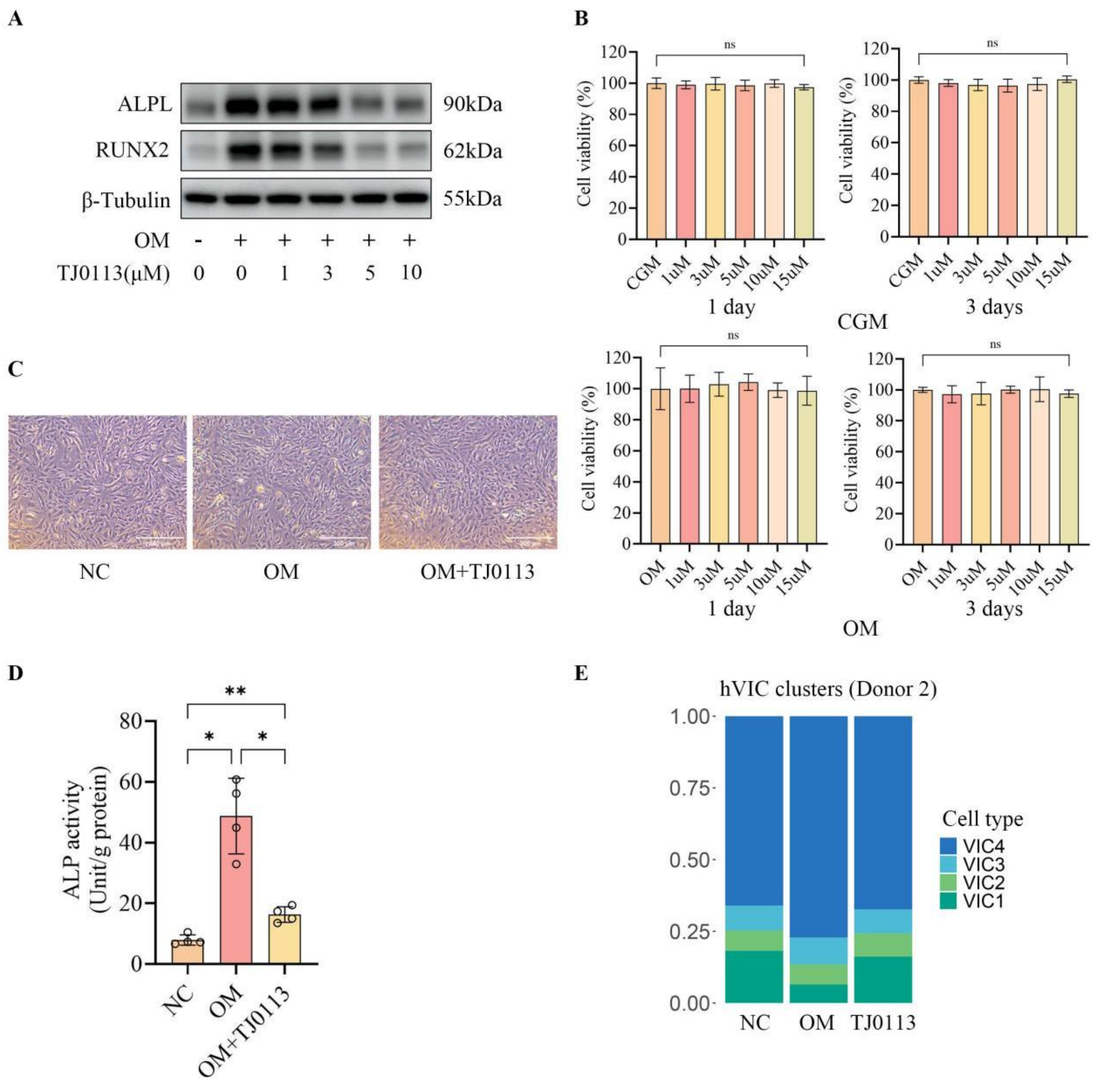


(A) Representative Western blot images of ALPL and RUNX2 in hVICs treated with different concentrations of TJ0113 under osteogenic conditions, identifying 5 μM as the effective concentration that reduced ALPL and RUNX2 expression. (B) CCK-8 assay showing that different concentrations of TJ0113 did not affect hVIC viability in complete growth medium (CGM, upper) or osteogenic medium (OM, lower) at 1 day (left) or 3 days (right) (n = 5). (C) Representative bright-field images showing that 5 μM TJ0113 did not alter hVIC morphology. (D) Alkaline phosphatase activity in the control, OM, and OM + TJ0113 groups, showing that TJ0113 reduced OM-induced ALP activity (n = 4). (E) scRNA-seq analysis of the proportions of each hVIC subcluster across groups in a second representative primary hVIC sample. Scale bars: (C) 300 μm. For the statistical analysis, *P < 0.05 and **P < 0.01 indicate significant differences, and ns indicates no significant difference.

**Supplementary Figure 3. TJ0113 does not affect global macroautophagy in hVICs.**

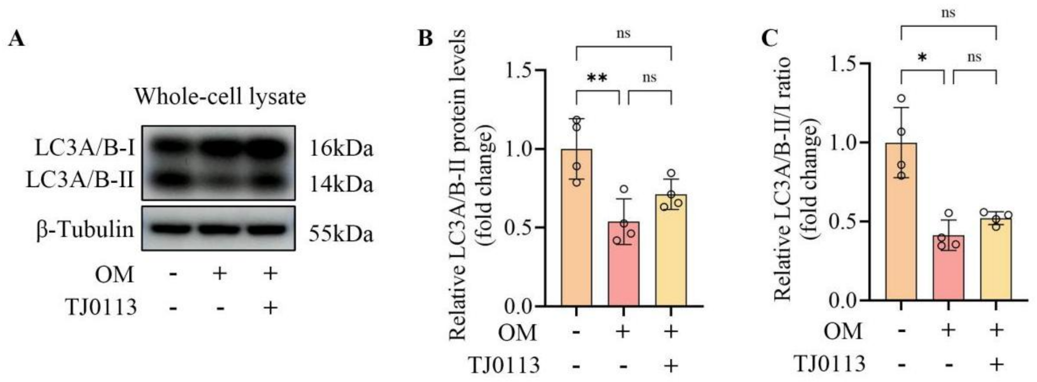


(A) Representative Western blot images of LC3A/B in whole-cell lysates of hVICs cultured in control medium, OM, or OM supplemented with TJ0113. (B) Quantification of LC3A/B-II levels (n = 4). (C) Quantification of the LC3A/B-II/I ratio (n = 4). For the statistical analysis, *P < 0.05 and **P < 0.01 indicate significant differences, and ns indicates no significant difference.

**Supplementary Figure 4. TJ0113 reduces ROS levels in hVICs under osteogenic conditions.**

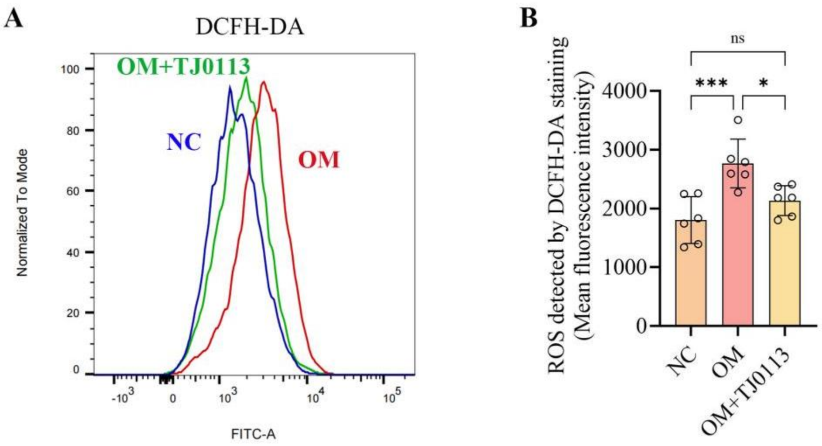


(A) Representative flow cytometry histograms of DCFH-DA fluorescence in hVICs cultured in control medium, OM, or OM supplemented with TJ0113. (B) Quantification of mean DCFH-DA fluorescence intensity (n = 6). For the statistical analysis, *P < 0.05 and ***P < 0.001 indicate significant differences, and ns indicates no significant difference.

**Supplementary Figure 5. PINK1 is upregulated upon osteogenic induction and its knockdown aggravates osteogenic differentiation.**

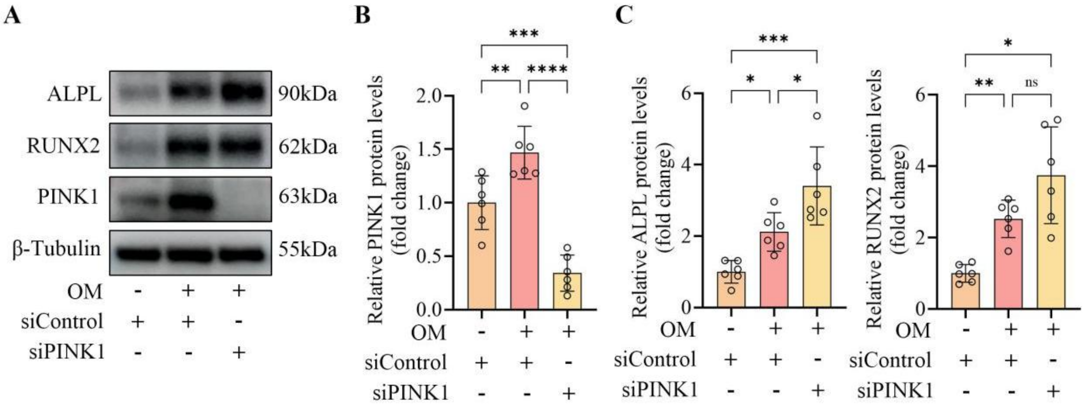


(A) Representative Western blot images of PINK1, ALPL, and RUNX2 in hVICs transfected with siPINK1 or siControl and cultured under osteogenic conditions. (B)

Quantification of PINK1 protein levels (n = 6). (C) Quantification of ALPL and RUNX2 protein levels (n = 6). For the statistical analysis, *P < 0.05, **P < 0.01, ***P < 0.001, and ****P < 0.0001 indicate significant differences, and ns indicates no significant difference.

**Supplementary Figure 6. Cardiac function and body weight in a high-fat diet-induced CAVD mouse model.**

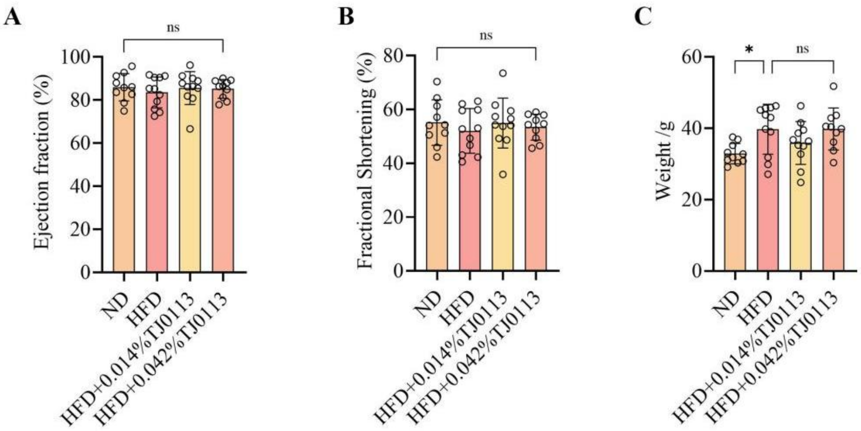


(A) Ejection fraction, (B) fractional shortening, and (C) body weight in the four groups: normal chow diet (ND), high-fat diet (HFD), HFD + 0.014% TJ0113, and HFD + 0.042% TJ0113 (n = 10–11). For the statistical analysis, *P < 0.05 indicates significant difference, and ns indicates no significant difference.

**Supplementary Figure 7. TJ0113 treatment does not affect plasma metabolic, hepatic, or renal parameters in a high-fat diet-induced CAVD mouse model.**

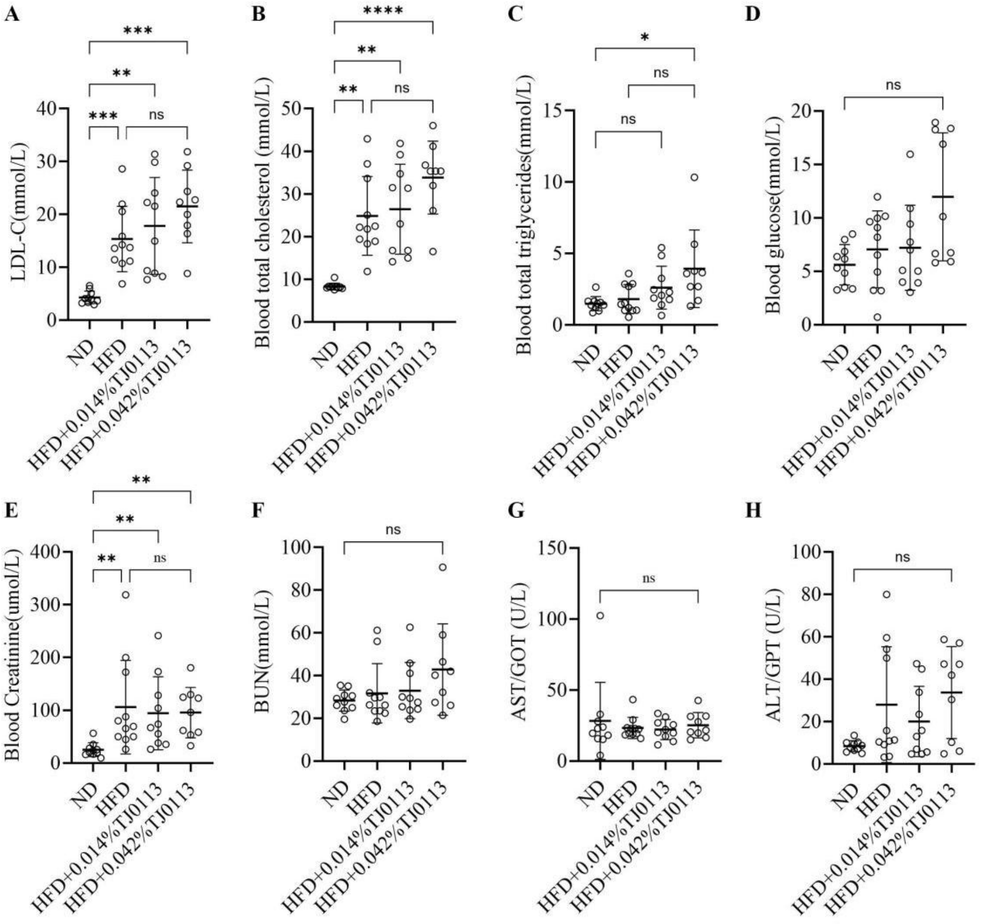


(A) LDL-C, (B) total cholesterol, (C) total triglycerides, (D) blood glucose, (E) creatinine, (F) BUN, (G) AST, and (H) ALT in the four groups: normal chow diet (ND), high-fat diet (HFD), HFD + 0.014% TJ0113, and HFD + 0.042% TJ0113 (n = 9–11). For the statistical analysis, *P < 0.05, **P < 0.01, ***P < 0.001, and ****P < 0.0001 indicate significant differences, and ns indicates no significant difference.

**Supplementary Figure 8. TJ0113 alleviates calcific aortic valve stenosis in a wire injury-induced CAVD mouse model.**

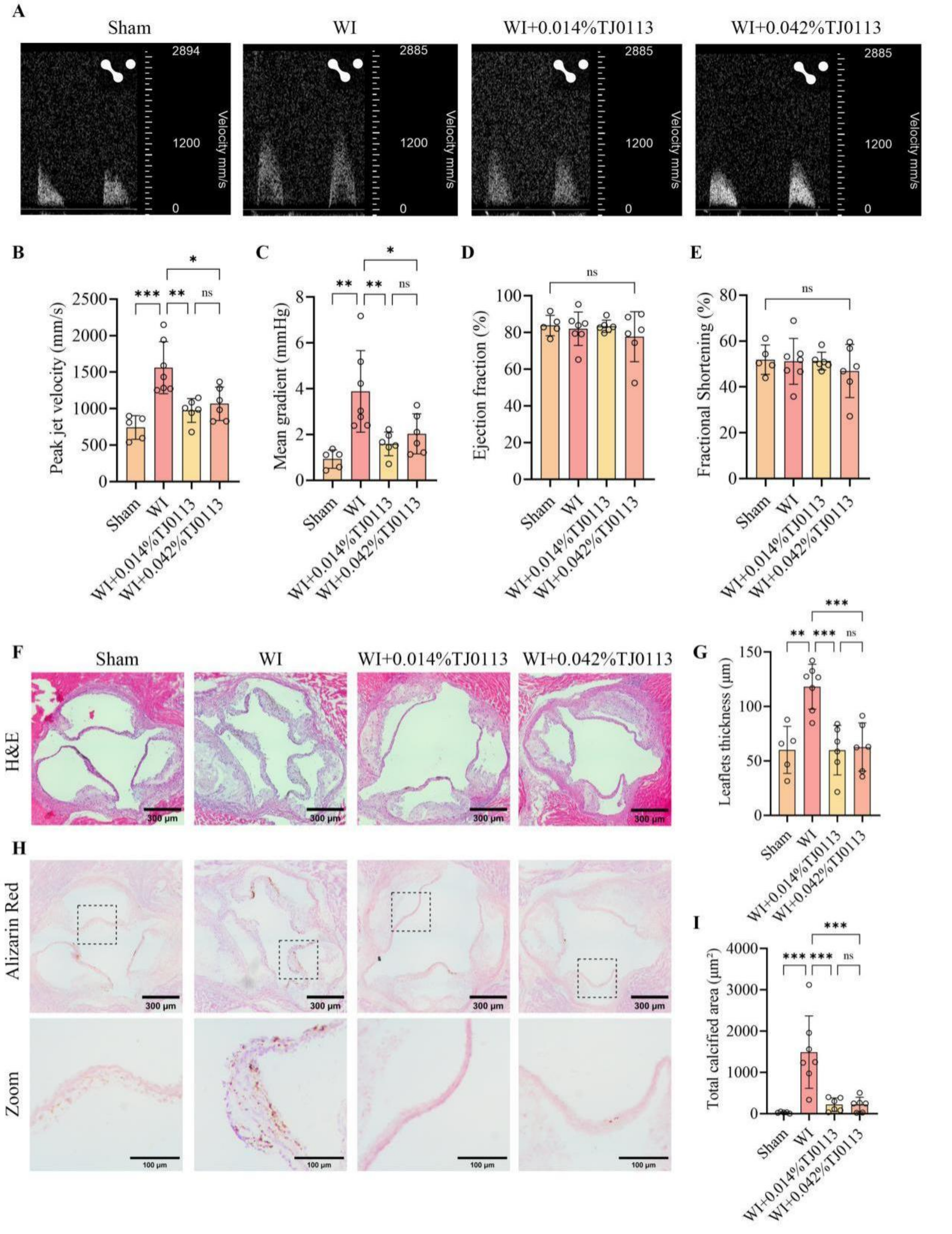


(A) Representative pulsed-wave Doppler images of aortic valve flow velocity in the four groups: Sham, wire injury (WI), WI + 0.014% TJ0113, and WI + 0.042% TJ0113. (B) Peak jet velocity. (C) Mean transvalvular pressure gradient. (D) Ejection fraction. (E) Fractional shortening. (F) Representative H&E staining images of mouse aortic valve leaflets. (G) Quantification of aortic valve leaflet thickness. (H) Representative Alizarin Red staining images of calcific nodule deposition in mouse aortic valves, with magnified views of boxed regions (lower panels). (I) Quantification of calcified area. n = 5–7 per group. For the statistical analysis, *P < 0.05, **P < 0.01, and ***P < 0.001 indicate significant differences, and ns indicates no significant difference. Scale bars: (F) 300 μm; (H) upper panels, 300 μm; lower panels (magnified views of boxed regions), 100 μm.

**Supplementary Figure 9. MCL1 and PRMT3 protein levels in hVICs cultured under different conditions.**

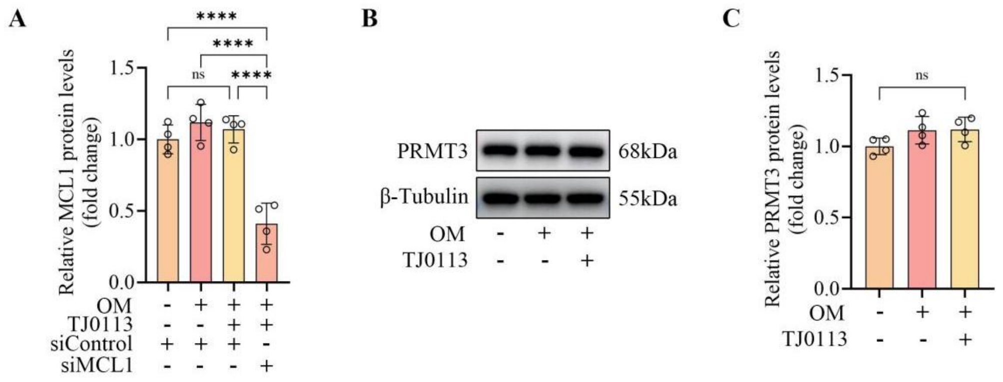


(A) Quantification of MCL1 protein levels in hVICs transfected with siMCL1 or siControl and cultured under different conditions (n = 4). (B) Representative Western blot images of PRMT3 in whole-cell lysates of hVICs cultured in control medium, OM, or OM supplemented with TJ0113. (C) Quantification of PRMT3 protein levels (n = 4). For the statistical analysis, ****P < 0.0001 indicates significant difference, and ns indicates no significant difference.

**Supplementary Figure 10. PRMT3 knockdown inhibits osteogenic differentiation of hVICs.**

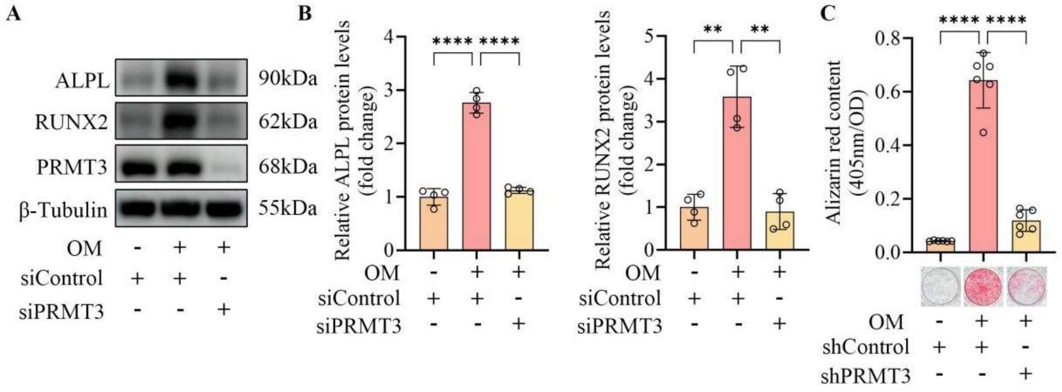


(A) Representative Western blot images of ALPL, RUNX2, and PRMT3 in hVICs transfected with siPRMT3 or siControl and cultured under different conditions. (B) Quantification of ALPL and RUNX2 protein levels (n = 4). (C) Alizarin Red staining and quantification of calcium deposition in hVICs transduced with shPRMT3 or shControl lentivirus and cultured under different conditions for 21 days (n = 6). For the statistical analysis, **P < 0.01 and ****P < 0.0001 indicate significant differences.

**Supplementary Figure 11. BACH1 protein levels remain unchanged across conditions in hVICs.**

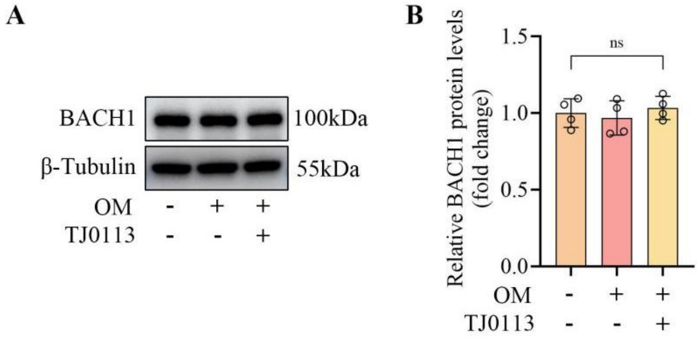


(A) Representative Western blot images of BACH1 in whole-cell lysates of hVICs cultured in control medium, OM, or OM supplemented with TJ0113. (B) Quantification of BACH1 protein levels (n = 4). For the statistical analysis, ns indicates no significant difference.

**Supplementary Figure 12. TJ0113 and PRMT3 knockdown reduce PARM1 protein levels in hVICs.**

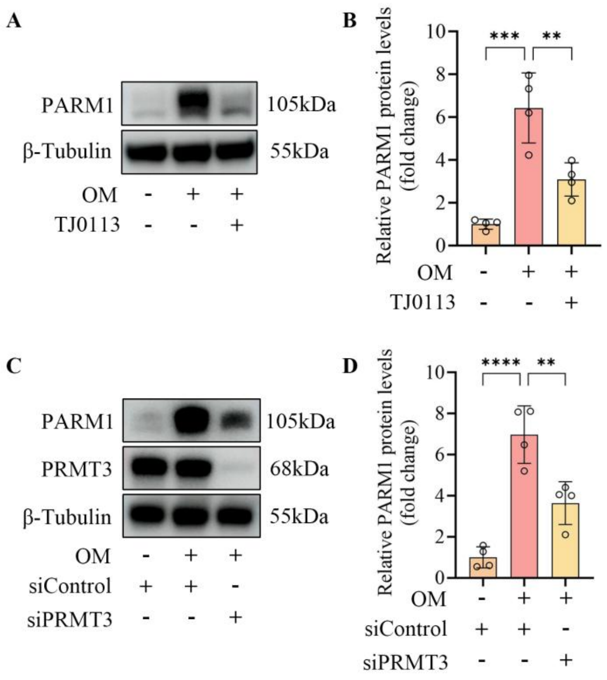


(A) Representative Western blot images of PARM1 in hVICs cultured in control medium, OM, or OM supplemented with TJ0113. (B) Quantification of PARM1 protein levels (n = 4). (C) Representative Western blot images of PARM1 in hVICs transfected with siPRMT3 or siControl and cultured under different conditions. (D) Quantification of PARM1 protein levels (n = 4). For the statistical analysis, $**P < 0.01$, $***P < 0.001$, and $****P < 0.0001$ indicate significant differences.

**Supplementary Figure 13. TJ0113 reduces PARM1 expression in aortic valves of wire injury-induced CAVD mice.**

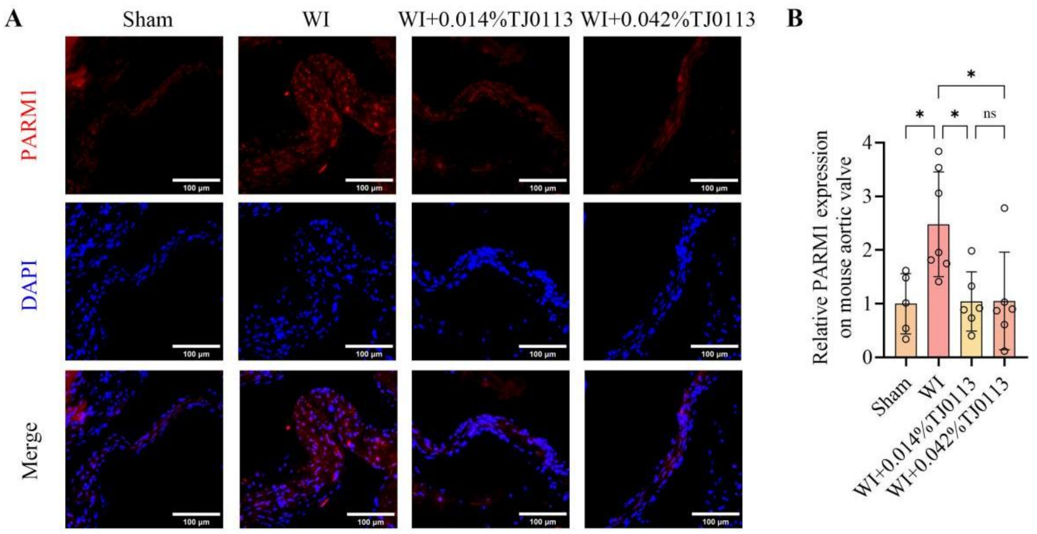


(A) Representative immunofluorescence images of PARM1 in aortic valve leaflets from the four groups: Sham, WI, WI + 0.014% TJ0113, and WI + 0.042% TJ0113; nuclei were counterstained with DAPI. (B) Quantification of PARM1 fluorescence intensity in mouse aortic valves (n = 5–7). For the statistical analysis, *P < 0.05 indicates significant difference, and ns indicates no significant difference. Scale bars: (A) 100 μm.

**Supplementary Table 1. Sequence for siRNA**

| siRNA name | | Sequence (5'-3') |
|---|---|---|
| siControl | sense | UUCUCCGAACGUGUCACGUTT |
| | antisense | ACGUGACACGUUCGGAGAATT |
| siPINK1 | sense | GCCAGUACCUUUGUGUGAATT |
| | antisense | UUCACACAAAGGUACUGGCTT |
| siMCL1 | sense | GUAGGGUGGAAAGACUUAATT |
| | antisense | UUAAGUCUUUCCACCCUACTT |
| siPRMT3 | sense | CCUUGGGAGAAAGAAGAGUAU |
| | antisense | AUACUCUUCUUUCUCCCAAGG |
| siBACH1 | sense | GUCAGGAUUUACCUUUGAATT |
| | antisense | UUCAAAGGUAAAUCCUGACTT |

**Supplementary Table 2. Primer sequences for RT-qPCR**

| Gene | | Sequence (5'-3') |
|---|---|---|
| hPARM1 | Forward | TCTTAACTGCAGGATGGAGGG |
| | Reverse | TAGTCCAGATGGTGGTCGGT |
| hTUBB | Forward | GGTTGATTCTGTCCTGGATGTGGTA |
| | Reverse | GGTATTCTTCTCGGATCTTGCTGATAAG |

**Supplementary Table 3. Primary antibodies**

| Antibodies | application | dilution | Supplier | Catalogue number |
|---|---|---|---|---|
| ALPL | WB | 1:1000 | R&D | MAB29092 |
| RUNX2 | WB | 1:1000 | CST | 8486 |
| HRP-β-Tubulin | WB | 1:10000 | Abclonal | AC030 |
| MCL1 | WB | 1:1000 | CST | 94296 |
| MCL1 | IP | 1:100 | CST | 94296 |
| LC3A/B | WB | 1:1000 | CST | 12741 |
| PINK1 | WB | 1:500 | HuaBio | HA723021 |
| PRMT3 | WB | 1:10000 | Abcam | ab191562 |
| PRMT3 | IP | 1:50 | Invitrogen | 703744 |
| VDAC | WB | 1:1000 | CST | 4661 |
| BACH1 | WB | 1:1000 | CST | 33059 |
| LaminB1 | WB | 1:10000 | Proteintech | 12987-1-AP |
| PARM1 | WB | 1:1000 | Abcam | ab168369 |
| PARM1 | IF | 1:200 | Biorbyt | orb100604 |
| Normal Rabbit IgG | IP | Matched by mass* | CST | 2729 |

*Normal Rabbit IgG was used as an IP negative control and was matched by mass (μg) to the primary antibody
